# The quantitative metabolome is shaped by abiotic constraints

**DOI:** 10.1101/2020.09.26.315135

**Authors:** Amir Akbari, James T. Yurkovich, Daniel C. Zielinski, Bernhard O. Palsson

## Abstract

Living systems formed and evolved under governing constraints that characterize their interactions with the inorganic world. These interactions are definable using basic physico-chemical principles. Here, we formulate a comprehensive set of ten governing abiotic constraints that define possible quantitative metabolomes. We apply these constraints to a metabolic network of *Escherichia coli* that represents 90% of its metabolome. We show that the quantitative metabolomes allowed by the abiotic constraints are consistent with metabolomic and isotope labeling data. We find that: (i) Network-wide characterization of charge-, proton- and magnesium-related constraints shape transcriptional regulatory responses to osmotic stress; (ii) Proton and charge imbalance underlie transcriptional regulatory responses to acid stress; (iii) Abiotic constraints drive the evolution of transport systems, such as high-affinity phosphate transporters. Thus, quantifying the constraints that the inorganic world imposes on living systems provides insights into their key characteristics, helps understand the outcomes of evolutionary adaptation, and should be considered as a fundamental part of theoretical biology and for understanding the constraints on evolution.

## 1. Introduction

Living organisms perform a variety of cellular functions, such as metabolism, protein synthesis, replication, cell division, and communication with the environment [1]. Systems-level descriptions of the operation, interconnection, and interaction of basic biochemical reaction networks provide a fundamental understanding of these essential life processes and their coordinated functions [2]. Driven by demand for explanatory and predictive frameworks, focus in systems biology has shifted towards comprehensive and mechanistic modeling approaches that can explain the functions of basic biological networks [3].

Constraint-based approaches to systems biology have proved successful [4, 5]. They integrate genome-scale reconstructions of metabolism [6] with regulatory-networks [7] and macromolecular-expression [8] networks. The latter simultaneously computes the activity states of the metabolic network and protein allocation needed to support and optimize cellular functions. Constraint-based approaches define a set of possible homeostatic states in a solution space, which is formed by stoichiometric, maximum-enzyme capacity, and other constraints.

One can search this space for homeostatic states of interest. Cellular objectives are used to define these homeostatic states. Constraint-based optimization is used to determine the best material and energy flow through the network to fulfill the stated objectives [9], and thus the detailed phenotype. The computed solutions provide the simultaneous activity of all the reactions in the network needed to reach a homeostatic state. Reaction activities are computed based on a flux balance that is independent of concentrations, which are thus not computed. Except for the direction of effectively irreversible reactions, no thermodynamic constraints are imposed.

In this article, we develop a constraint-based approach that is focused on metabolite abundances. Possible concentration ranges are estimated by identifying quantitative metabolomes that satisfy all fundamental and evolutionary physico-chemical constraints imposed on biological networks. A total of ten abiotic constraints (ABCs) are identified as governing allowable metabolite concentrations that have to be satisfied simultaneously to form a space of all possible quantitative metabolomes. The ABCs are universal and should apply from primitive to modern cells across the tree of life. We show that the ABCs shape transcriptional regulation, transporter functions, and are consistent with experimental measurements.

## 2. Results

### 2.1. Formulation of ABC-based analysis

#### Stating the ABCs

We introduce an ABC-based approach to characterize the constraints that govern the interactions of biological functions with the inorganic world. We consider ten fundamental and evolutionary classes of abiotic constraints on cellular functions (Fig. 1 and Abiotic Constraints): (i) thermodynamics, (ii) charge balance, (iii) solubility, (iv) membrane potential, (v) buffer capacity, (vi) enzyme saturation, (vii) ionic strength, (viii) osmotic balance, (ix) ion binding, and (x) unresolved metabolites (see Fig. 1A).

**Figure 1:**
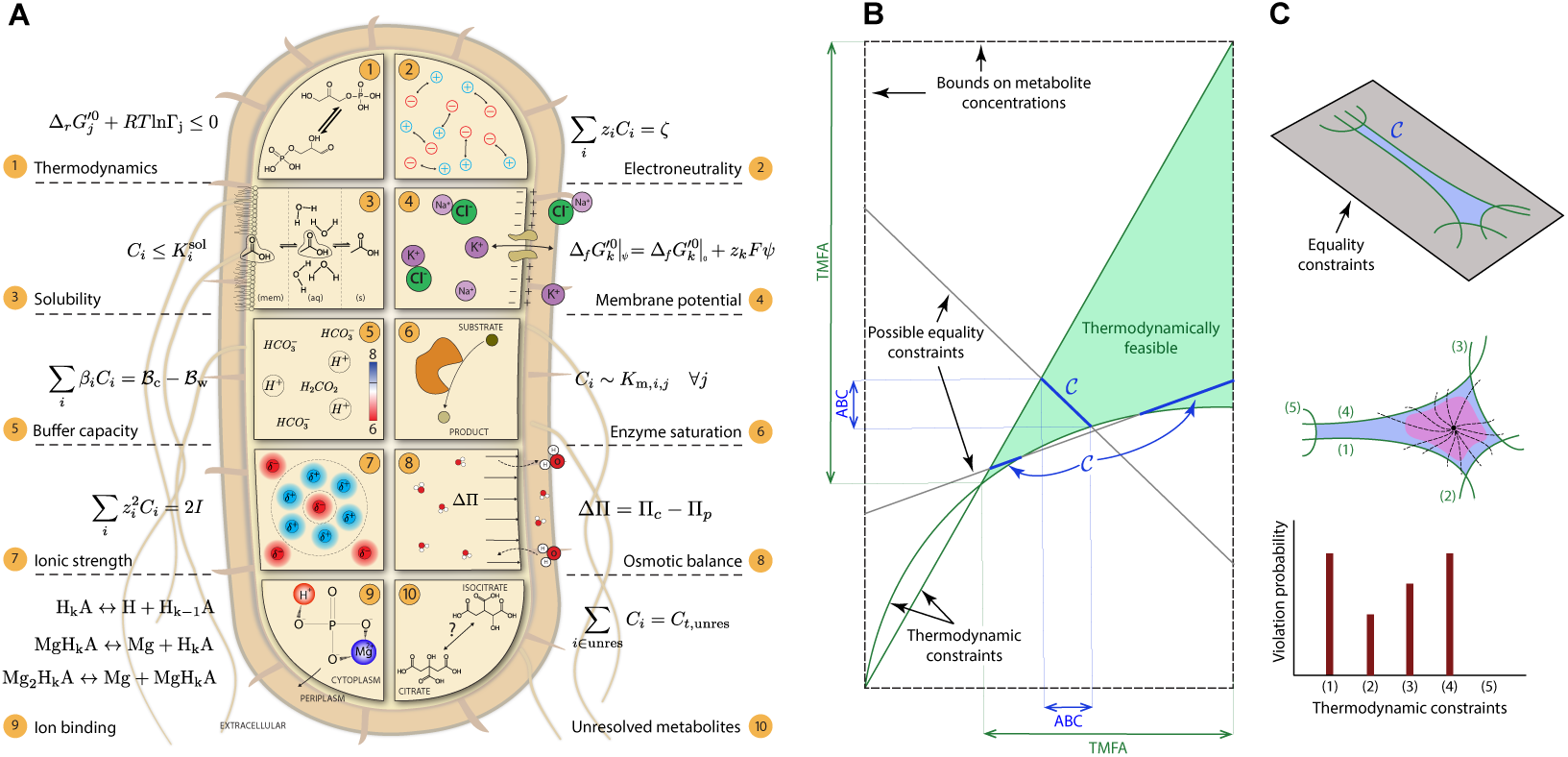
A constraint-based approach to determine the concentration state of metabolic networks. (A) Schematic representation of ten fundamental and evolutionary abiotic constraints that together define a concentration solution space 𝒞. Reactants, reactions, and species are associated with the indices *i, j*, and *k*, respectively. (B) Construction of 𝒞 from the ABCs, formulated as equalities (*e.g*., charge balance) and inequalities (*e.g*., thermodynamic), illustrated in two-dimensions. The solution space 𝒞 can be convex or disconnected. Feasible concentration ranges furnished by the ABCs are generally more restricted than those predicted by thermodynamic constrains alone (TMFA analysis [10]). (C) Sampling the solution space by generating random trajectories in 𝒞 from an interior point. Interior points are identified in a region (purple area), where experimental data most likely fall. Trajectories are continued until one of the thermodynamic constraints is violated. For each thermodynamic constraint, the violation probability is defined as the ratio of the number of trajectories intersecting it and the total number of trajectories.

Mathematically, thermodynamic and solubility constraints are represented by inequalities and the rest by equalities. Together, these ten classes of constraints define a nonconvex and disconnected set of possible concentration states that is challenging to characterize (see Fig. 1B). The equality constraints couple all the thermodynamic constraints together, so that metabolite concentrations can be restricted by the thermodynamic constraints of any reaction in the network, regardless of the associations between reactions and metabolites. Throughout the paper, we refer to the ABC-based analysis as the ABC for brevity.

#### The conceptual basis

The ABC is analogous to the widely-used flux-balance analysis (FBA) [6]: A biologically relevant concentration solution space (CSS) is constructed by restricting the concentrations to those obeying all the abiotic constraints. Cellular objectives are evaluated according to their predictive accuracy by determining associations between the optima of the objective functions and intracellular concentration data. These objectives play the same role in the ABC as the growth rate does in FBA. The mathematical formulation and computational approaches are described in STAR Methods.

#### Model system used for analysis

We applied the ABCs to characterize the CSS of a reduced metabolic network of *E. coli*, comprising 78 reactions, 72 cytoplasmic, and 11 periplasmic metabolites (Fig. S3). This network consists of: (i) high-flux pathways and (ii) pathways connecting the reduced metabolism to key metabolites with the largest concentrations among those reported in the literature [11, 12]. Major pathways, such as glycolysis, tricarboxylic acid (TCA) cycle, anaplerotic reactions, electron transport chain (ETC), biosynthetic sugar metabolism, and cofactor interconversions are included in this network. Importantly, this network contains over 90% of the observed metabolome by mole (note that moles, not mass, determine osmotic pressure) and includes all reactions with experimentally measured fluxes [12]. Therefore, it possesses the essential characteristics of the global metabolome for the ABC.

#### Graphical representation of results

The computation of the ABCs provides the role that each metabolite and each reaction plays in satisfying the ABCs. These different roles, which can be displayed graphically, include: (i) metabolite concentration and reaction energy, (ii) charge distribution, (iii) proton distribution, and (iv) magnesium distribution (See Figure 3). These graphs show various ways, in which biochemical reactions can influence the homeostatic states of the reduced network. For example, panels (B)-(D) highlight the reactions and metabolites that are the major players in charge, proton, and magnesium homeostasis. Therefore, they are parts of the network that can most effectively counterbalance stress-induced perturbations in these three quantities. Overall, these maps visualize the systems biology of stress responses.

#### Synopsis of the results

We present our results in three parts. First, we show how the ABCs can interpret and predict concentration and flux data. We also elaborate on the implications of our analysis for other constraint-based approaches, such as thermodynamics-based metabolic flux analysis (TMFA) [10]. Then, we provide two cases, illustrating how the ABCs shape transcriptional responses to stress conditions. Finally, we show that the ABCs necessitate the evolution of multiple specialized transporters for phosphate.

### 2.2. Feasible concentration ranges defined by the ABCs are consistent with metabolomic data

#### Metabolomic data

We sought to determine whether the ABCs were consistent with available experimental metabolomic data. We considered growth on four carbon sources, including glucose, acetate, pyruvate, and succinate. The flux states of the network were determined using the latest genome-scale metabolic network reconstruction of *E. coli* [13]. The FBA model (Flux Distribution; STAR Methods) was solved to identify metabolic fluxes that best matched their experimentally measured values [12] for each of the four carbon sources. The resulting flux directions were then used to specify the thermodynamic constraints for the ABC.

#### Computing the consistency of metabolomic data with the ABCs

We identified the theoretical cytoplasmic concentrations *C*_th_ (Fig. 2B, crosses), that satisfied all the ABCs and best matched the measured concentrations *C*_ex_ [12] (Fig. 2B, circles) for each carbon source. We found a strong correlation (corr(*C*_th_, *C*_ex_) = 0.84–0.93) and relatively small root-mean-square deviation (RMSD = 2.3–5.1 mM) between the computed and experimental concentrations across all four carbon sources (Fig. 2A), showing that the CSS contains the measured physiologically-relevant concentration states.

**Figure 2:**
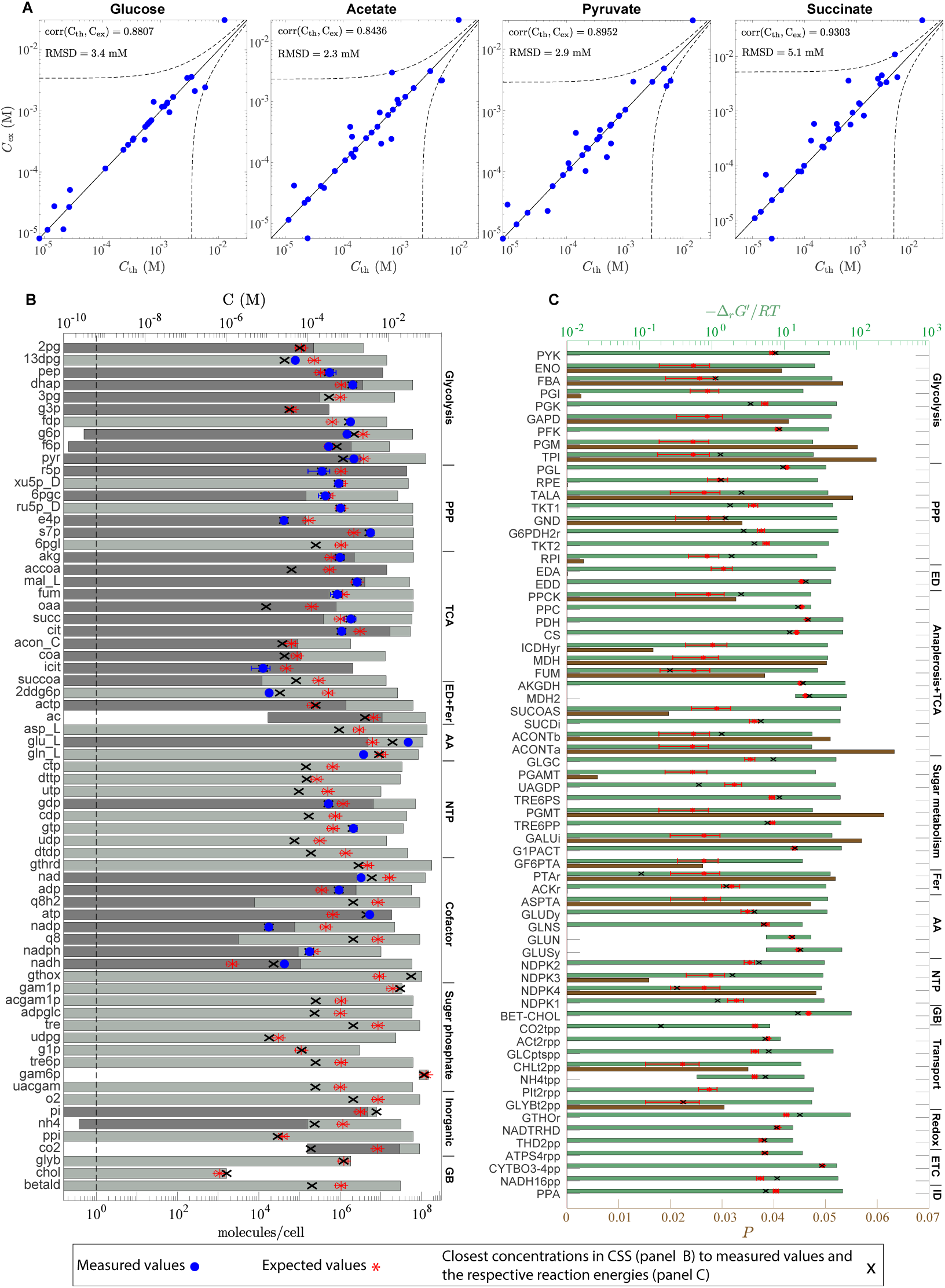
The ABCs are consistent with metabolomic and isotope-labeling data for *E. coli*. (A) Pearson correlation coefficient of measured concentrations *C*_ex_ and the closest theoretical concentrations *C*_th_ satisfying all the ABCs. Dashed lines indicate deviation by RMSD from the line *C*_ex_ = *C*_th_. (B) Feasible ranges of metabolite concentrations. Dark gray bars, determined from known Michaelis constants [14] (Eqs. (53) and (54); STAR Methods), show part of the feasible range, where some of the reactions a given metabolite participates in are undersaturated. Measured concentrations are reported by Gerosa et al. [12]. The number of molecules per cell is calculated from the molar concentration based on *V*_cell_ = 2.6 fL [15]. (C) Feasible ranges of reaction Gibbs energies and violation probabilities of thermodynamic constraints. For (B) and (C), glucose is the sole carbon source. Pathway abbreviations are defined in Table S3. Error bars indicate twice the standard deviation.

**Figure 3:**
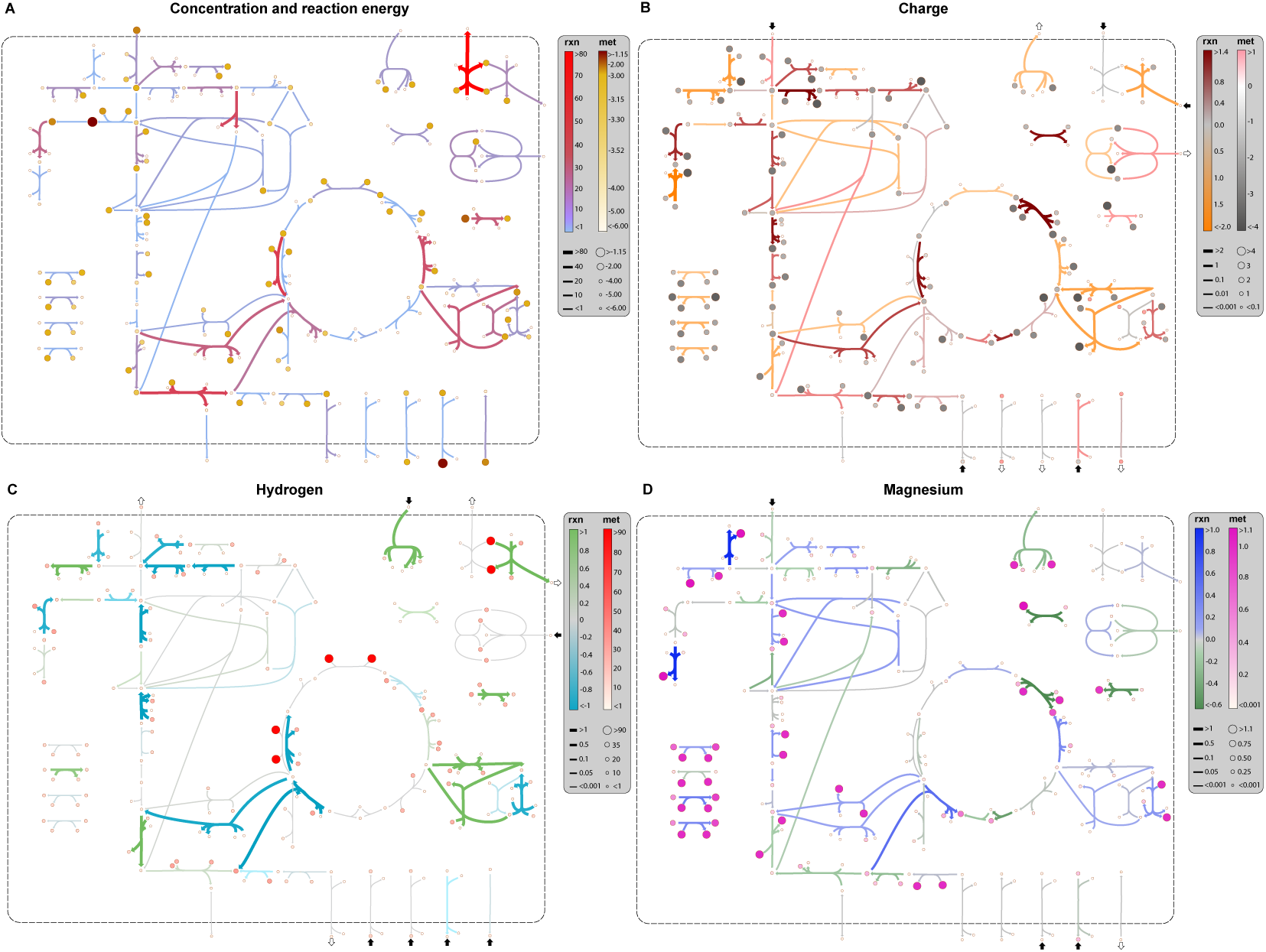
Application of the ABCs reveal the biophysical mechanisms underpinning the *E. coli* metabolic network. Computations are performed for exponential growth on glucose under physiological conditions (pH_*p*_ = 7, pH_*c*_ = 7.5, [Mg]_*c*_ = 2 mM, *I*_*c*_ = 300 mM). (A) Concentration and energy map: reaction colorbar indicates reaction Gibbs energy in kJ/mol-rxn and metabolite colorbar indicates log_10_(*C*_*i*_*/C*^*º*^) with *C*^*º*^ = 1 M. (B) Charge map: reaction colorbar indicates intrinsic charge consumption in mol-*e*/mol-rxn and metabolite colorbar indicates metabolite effective-charge in mol-*e*/mol-met. (C) Hydrogen map: reaction colorbar indicates intrinsic proton consumption in mol-H/mol-rxn and metabolite colorbar indicates metabolite hydrogen content in mol-H/mol-met. (D) Magnesium map: reaction colorbar indicates intrinsic magnesium consumption in mol-Mg/mol-rxn and metabolite colorbar indicates metabolite magnesium content in mol-Mg/mol-met. Metabolite and reaction colorbars are denoted by ‘met’ and ‘rxn’, respectively. For (B)-(D), arrows point in the direction, in which negative charge, proton, and magnesium ions are consumed. Reaction charge-, proton-, and magnesium-consumption for transport reactions, as indicated by the colorbars, do not reflect translocated protons. Small arrows next to each transport reaction show the direction of their respective net fluxes into and out of the cell, accounting for translocated protons and the accompanied metabolites. Enlarged and detailed maps are provided in Figs. S7– S10. See Abiotic Constraints for the definition of reaction charge-, proton-, and magnesium-consumption.

#### Highly connected metabolites in the network have more restricted concentration ranges

We computed feasible concentration ranges for all intracellular metabolites in the reduced network when glucose is the sole carbon source (Fig. 2B). The ABC reveals a qualitative connection between the network topology and the global concentration bounds. The upper and lower concentration bounds are generally restrictive for hub metabolites (*e.g*., glucose 6-phosphate and fructose 6-phosphate), while they exhibit the opposite trend for end-node metabolites (*e.g*., trehalose and UDP-N-acetyl-glucosamine). The hub metabolites simultaneously participate in several reactions, and their possible concentrations are constrained to a narrow range, so that they satisfy all the thermodynamic constraints placed on these reactions. The lower bound for exported metabolites with a low periplasmic concentration is restrictive (*e.g*., CO_2_). This restriction is due to the fact that the cell must retain a minimum cytoplasmic concentration to sustain a thermodynamic driving force across the cell membrane. For a similar reason, the upper bound for imported metabolites with a low periplasmic concentration is also restrictive (*e.g*., choline).

#### Computation of expected values of metabolite concentrations

To derive order-of-magnitude estimates of cytoplasmic concentrations, we computed expected values for concentrations (Fig. 2B, asterisks) by sampling allowable metabolome compositions in the interior of the CSS (Characterization of Concentration Solution Space; STAR Methods). We found a good agreement between the expected values and measured concentrations for all metabolites, except for a few cofactors, such as ATP, NADP, and NADH (RMSD = 5.47 mM, excluding the cofactors). These cofactors participate in many reactions across the network. They are highly regulated with concentrations spanning several orders of magnitude depending the growth condition or in response to environmental stresses [10, 12]. As a result, their concentrations may be driven towards the CSS boundaries to accommodate a desired flux state. Thus, they cannot be generally well-approximated by the expected values, which are usually interior points of the CSS.

#### Comparing measured concentrations to Michaelis constants

Enzyme-saturation efficiency has been proposed in the literature as a cellular objective to determine optimal intracellular concentrations [16]. To maximize this objective, metabolite concentrations are expected to be of the same order as, or greater than, the Michaelis constants of the reactions the metabolites participate in. However, this heuristic principle does not apply to all enzymes (*e.g*., degradation reactions) [11]. Nevertheless, it is expected to hold for reactions in central-carbon metabolism, where flux directions alter in response to stress conditions, so that enzymes are efficiently saturated in both directions [11].

To examine this objective, we compared the maximum Michaelis constant 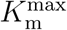 (Eq. (53); STAR Methods) of metabolites (Fig. 2B, dark gray bars) with measured concentrations (Fig. 2B, circles). We performed this comparison only for reactions with known kinetic parameters. We found 14 undersaturated reactions 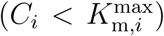 from central-carbon pathways (*e.g*., PGI, ICDHyr, TALA, and AKGDH), indicating that enzyme efficiency is not a fundamental cellular objective or constraint, with which to determine the concentration state of metabolic networks. This result is consistent with previous studies on the efficiency of enzymatic reactions [11, 17].

#### The ABCs yield more restrictive concentration bounds

The inclusion of charge-related constraints in the ABC limits the intracellular concentrations to biologically relevant ranges more restrictively than TMFA. We computed the concentration bounds arising from the ABCs and found narrower feasible concentration ranges than those predicted by TMFA, which only applies the thermodynamic constraints. (Fig. 5A).

**Figure 4:**
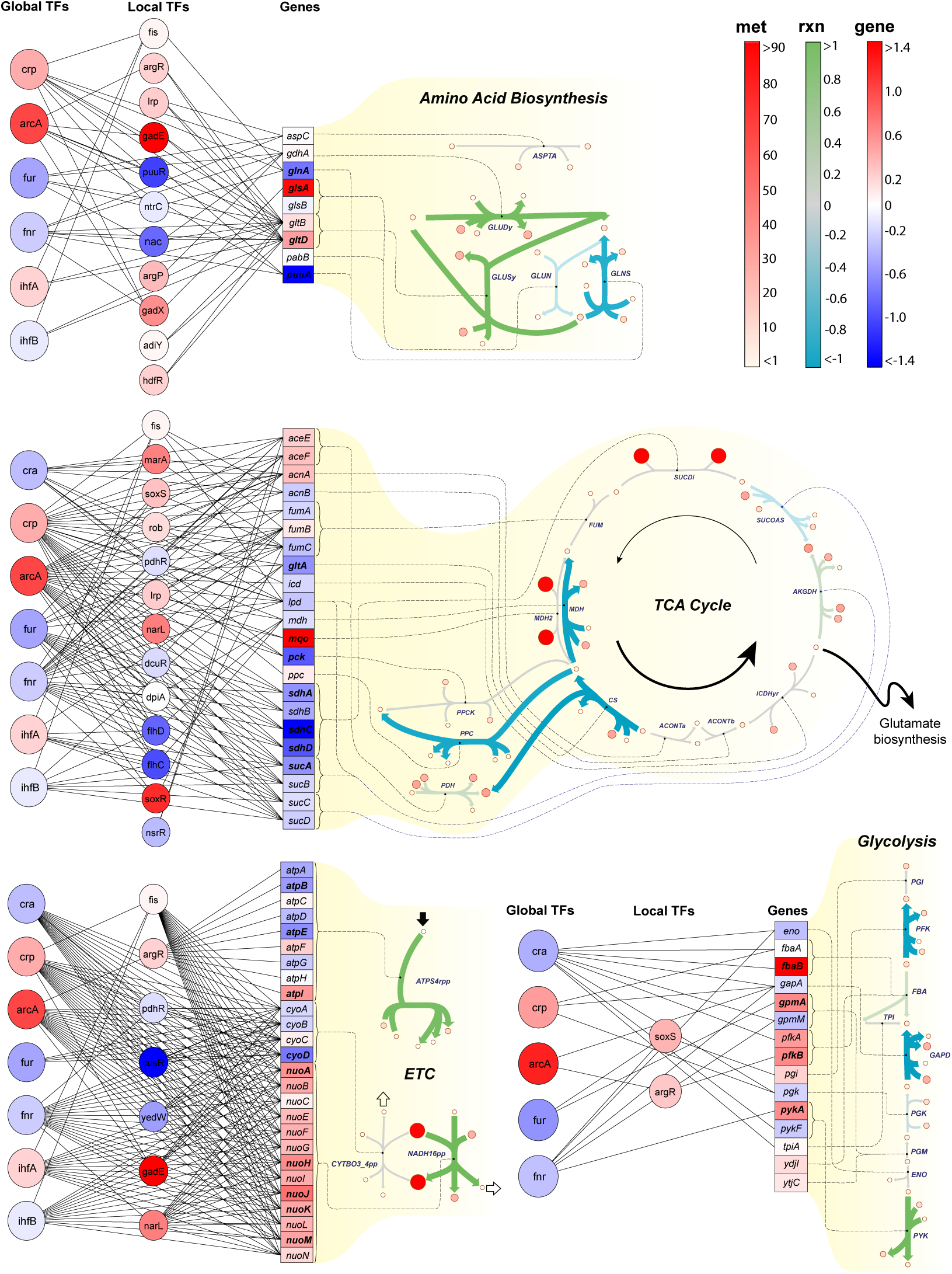
The association of transcriptional regulatory responses with key metabolic pathways reveals influences on the *E. coli* acid-stress response (pH_*p,ϵ*_ = 5.5). Reaction colorbar (rxn) indicates proton consumption in mol-H/mol-rxn, metabolite colorbar indicates metabolite hydrogen content in mol-H/mol-met, and gene-expression colorbar (gene) indicates expression change log_2_(TPM_*ϵ*_*/*TPM_0_). The subscripts *E* and 0 denote the acid-stress condition and base state. The base state corresponds to a stress-free growth in a neutral medium with pH_*p*,0_ = 7, pH_*c*,0_ = 7.5, [Mg]_*c*,0_ = 2 mM, and *I*_*c*,0_ = 300 mM. Genes with more than 1.4-fold expression change (shown in boldface) are considered differentially expressed. Expression data are taken from the RNA-sequencing analysis of Seo et al. [32] on a wild-type *E. coli* strain. Reaction proton-consumption for the ETC reactions, as indicated by the colorbar, does not reflect translocated protons. Small arrows next to each ETC reaction show the direction of net proton flux into (black) and out of (white) the cell.

**Figure 5:**
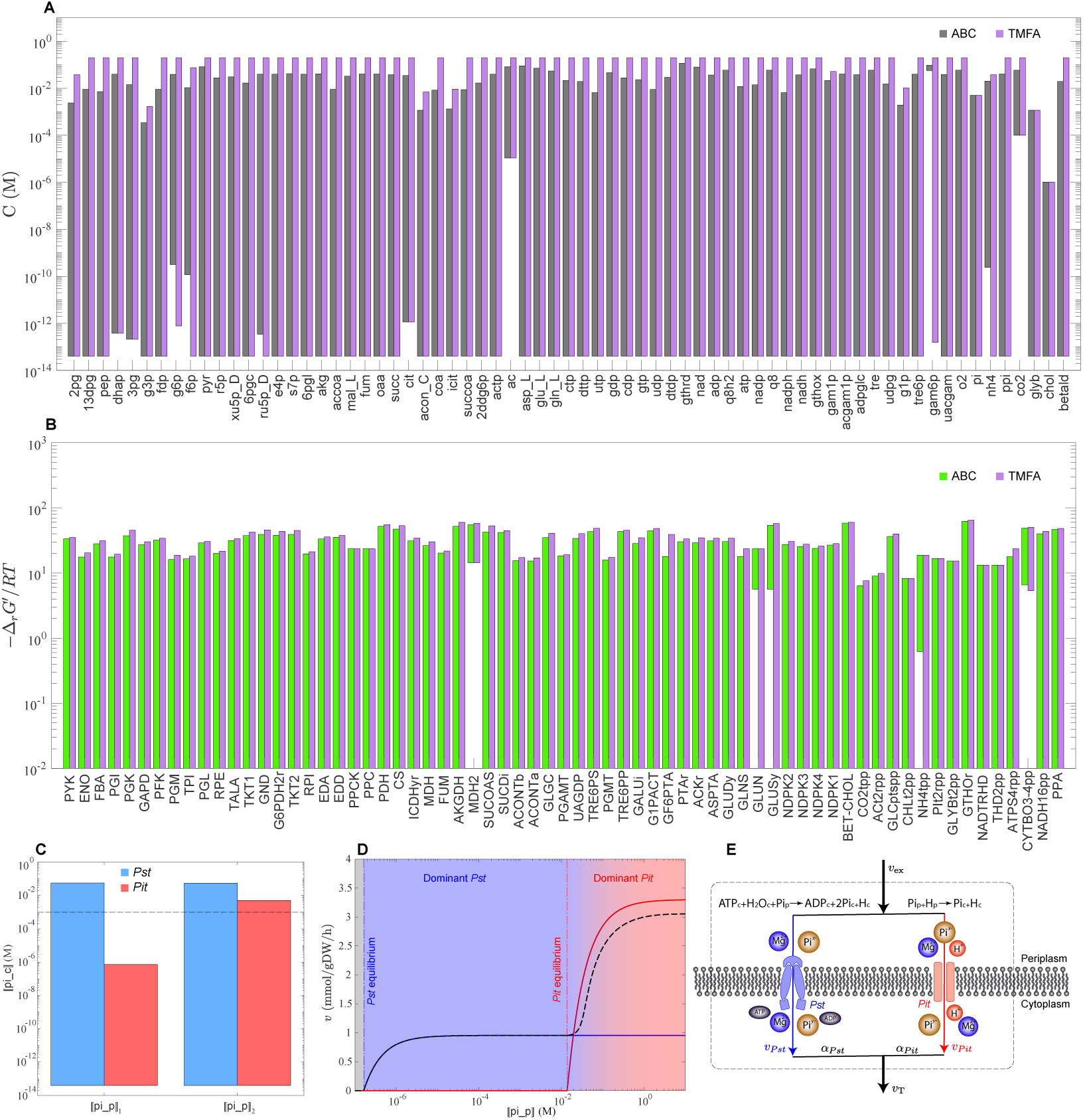
Abiotic constraints drive the evolution of alternative transport systems. Feasible ranges of metabolite concentrations (A) and reaction Gibbs energies (B) furnished by the ABC and TMFA are compared. (C) Feasible cytoplasmic-phosphate concentration ranges in mutants, where either the *Pst* system (blue) or the *Pit* system (red) is active at periplasmic concentrations ⟦pi_p⟧ _1_ = 0.01 and ⟦pi_p⟧ _2_ = 69 mM. (D) Phosphate-transport kinetics in *E. coli*. Solid blue and red lines show phosphate uptake rates in mutants, where either the *Pst* system or the *Pit* system is active, and dashed black line corresponds to a wild-type strain, where both systems are active. Maximum velocities and Michaelis constants are taken from Willsky and Malamy [36]. The total phosphate uptake rate is estimated *v*_T_ = *α*_*Pst*_*v*_*P st*_ + *α*_*Pit*_*v*_*Pit*_ with *α*_*Pst*_ = 0.09 and *α*_*Pit*_ = 0.9 at large ⟦pi_p⟧ [36]. Uptake rates are estimated using ⟦pi_c⟧ = 1, ⟦atp_c⟧ = 3.45, and ⟦adp_c⟧ = 0.6 mM. (E) Schematic representation of the *Pst* and *Pit* systems operating in parallel in wild-type strains.

#### Recap

The results of the ABC are consistent with high quality metabolomic data and other known network features. By comparing metabolite concentrations and Michaelis constants involved in the central-carbon metabolism of *E. coli*, we found that enzyme efficiency may not be a fundamental cellular objective.

### 2.3. Thermodynamic bottlenecks furnished by the ABC are consistent with isotope-labeling measurements

#### Thermodynamics play a fundamental role in adaptation

Modern bacteria have an extraordinary ability to grow and survive using minimal environmental resources. To maximize their survival chances, they efficiently allocate available resources to essential biochemical reactions, maintaining a steady flow of materials through all their metabolic pathways. Sustaining this optimal flux state hinges on a trade-off between pathway energetics and proteome allocation. Reactions with a small thermodynamic driving force have higher protein cost and vice versa [18]. Through this tradeoff between the thermodynamic driving force and proteome allocation, metabolic fluxes adapt to environmental perturbations [19].

#### Determining thermodynamic bottlenecks

We, thus, sought to determine the thermodynamic bottlenecks resulting from the ABCs. We calculated the feasible ranges of reaction Gibbs-energies (Fig. 2C, green bars) and the violation probabilities (Eq. (92); STAR Methods) of their respective thermodynamic constraints (Fig. 2C, brown bars) to identify the most constraining reactions of the reduced network when glucose is the sole carbon source. Reactions with negative upper and lower Gibbs-energy bounds (*e.g*., MDH2) are irreversible; all the other reactions can be either inactive or reversible. In FBA, every reaction is considered reversible if no information about Gibbs energies is available. However, using the feasible Gibbs-energy ranges furnished by the ABC, the flux solution space can be reduced by imposing the corresponding flux bounds. We also found that the violation probability of reactions with the Gibbs-energy expectation 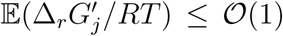 is substantially larger than other reactions. This serves as a qualitative criterion for when a reaction is thermodynamically constraining in a metabolic network.

#### Graphical representation of bottlenecks

We highlighted the thermodynamic bottlenecks of the reduced network on a network map (Fig. 3A) that visualizes metabolite-concentration distribution and reaction energies. These were evaluated at a point inside the CSS with the minimum distance from the experimental data of Gerosa et al. [12]. We found several reactions in glycolysis (ENO, PGI, GAPD, PGM), TCA-cycle (ICDHyr, MDH, ACONTa, SUCOAS), and sugar-metabolism (PGMT, PGAMT, GF6PTA, GALUi) pathways with small reaction Gibbs energies (|Δ_*r*_*G*^′^| ≪ 1). The feasible metabolite concentration ranges participating in these reactions are not necessarily restrictive, so they are regulated to operate close to their equilibrium.

#### Putting the results in context

The foregoing results can be contrasted to other studies in the literature. First, they are consistent with previous reports based on ^2^H and ^13^C metabolic flux analysis for glycolysis [19] and TCA cycle [20, 21]. Second, flux-direction reversal in glycolysis is often induced by carbon-source and nutrient alterations [12]. Thus, having near-equilibrium steps enhances the energy efficiency of the pathway and enables the cell to rapidly switch flux directions in response to energy and biomass demands with minimal concentration changes [19]. Third, the analysis indicates that there are several thermodynamic bottlenecks in the TCA cycle, hampering its full cyclic operation [20]. Fourth, the results suggest that thermodynamic efficiency (Enzyme Kinetics; STAR Methods) is not a fundamental cellular objective that determines the concentration state of metabolic networks.

#### The ABCs provide more restrictive bounds on reaction energies than thermodynamic constraints alone

Global concentration bounds derived from the ABCs can either coincide with (*e.g*., upper bound for choline) or be more restrictive than (*e.g*., lower bound for glucose 6-phosphate) those predicted by TMFA. The thermodynamic constraints are more dominant in the first case, and charge-related constraints are more dominant in the second. The dominant constraints set the limit for the metabolite concentrations. We compared the feasible ranges of reaction Gibbs energies arsing from the ABC and TMFA (Fig. 5B), identifying five irreversible reactions (GLUN, GLUSy, MDH2, NH4tpp, CYTBO3-4pp) compared to the three predicted by TMFA (MDH2, NH4tpp, CYTBO3-4pp).

#### Recap

The ABC is consistent with and unifies a broad range of disparate studies in the literature. We identified several near-equilibrium reactions in the central-carbon metabolism of *E. coli*, showing that thermodynamic efficiency is not a fundamental cellular objective.

### 2.4. Abiotic constraints shape transcriptional regulatory responses to environmental stresses

#### Ion-binding and charge-balance constraints lead to fundamental regulatory challenges

Biochemical reactions comprise transformations among metabolites that can have multiple charge states arising from proton dissociation and metal-ion binding [22]. Such reactions would liberate or bind protons and metal ions and, thus, interact with the cytoplasmic fluid to establish intracellular pH and metal-ion homeostasis. The reactions having the highest rate of exchange with the cytoplasmic fluid (highlighted in Figure 3) represent the key players in maintaining proton and metal-ion concentrations. Thus, they are prime targets for transcriptional regulations in response to environmental stresses.

#### Identifying major contributors to achieving charge-balance

Under physiological conditions, there is a balance between reactions donating negative charge to (Fig. 3B, orange arrows) and accepting negative charge from (Fig. 3B, brown arrows) the cytoplasmic fluid, that furnishes a stable electroneutral buffer for cellular metabolism. For example, in response to osmotic stress, *E. coli* imports osmoprotectants (*e.g*., glycine betaine and trehalose) or K^+^ from the extracellular environment to modulate its intracellular osmolarity, keeping a fixed osmotic pressure across the cell membrane [23]. The imbalance caused by these charged solutes can be alleviated by upregulating reactions producing counterions or downregulating those consuming counterions.

#### Known stress-response mechanisms

To maintain electroneutrality, the cell has many degrees of freedom, as to which reactions to upregulate and which ones to downregulate. Importantly, the robustness of *E. coli* to various environmental stresses suggests that the transcriptional regulatory network (TRN) performing osmoregulation has evolved to sustain homeostasis with respect to other simultaneous disturbances (*e.g*., carbon-source changes and pH stress), minimally impacting pathways performing other critical cellular functions.

Many genes in the *E. coli* genome are involved in osmoregulation. Primary transcriptional regulatory targets include potassium transporters (*e.g*., *trkA* and *kdpA*) [23, 24], osmoprotectant transporters (*e.g*., *proV*) [24], carbon-source transporters (*e.g*., *crr* and *ptsG*) [25], sodium transporters (*e.g*., *nhaA* and *nhaB*) [26], glutamate transporter (*e.g*., *gadC* and *gltI*) [27], and the ETC (*e.g*., *atpE* and *cyoD*) [28]. Secondary responses affect several intracellular pathways, such as amino-acid biosynthesis, osmoprotectant biosynthesis, central carbon metabolism, and nucleotide biosynthesis [25].

#### ABC-based interpretation of observed transcriptional responses

Short-term transcriptional regulatory responses to acid and osmotic stress are usually classified into phase I (≲ 20 min; regulation is transient) and phase II (20– 60 min; new steady state is reached) [26]. Early stress responses by wildtype strains differ from those exhibited by evolved strains that have adapted to high osmolarities [24]. In our formulation, strain-specific characteristics are captured by the active pathways, dominant metabolite concentrations, and condition-specific parameters of the reduced network. We constructed a charge map (Fig. 3B) using the characteristics of a wild-type *E. coli* strain to interpret short-term stress responses. Accordingly, we sought to establish associations between phase-I/II osmoregulatory responses and reaction charge-consumption provided by the ABC.

From the RNA-sequencing data of Seo et al. [27], we identified 69 differentially expressed genes (*>* 2-fold change 30 min after osmotic upshift) out of 124 genes associated with the reduced network. Among the primary-response genes that are differentially expressed, those associated with the ETC are all downregulated and those with osmoprotectant transporters are upregulated. Although not explicitly included in the reduced network, potassium importers and sodium exporters are also activated. Moreover, several intracellular reactions are among the secondary regulatory targets. For example, glycolytic reactions (except GAPD) and glutamate decarboxylase (GLUDC) (not explicitly included in the reduced network) are activated, while glutamate synthase (GLUSy) and glutamate dehydrogenase (GLUDy) are suppressed.

#### Deciphering regulatory principles

The foregoing observations signify the fundamental principles underlying the evolution of TRNs in bacteria to handle osmotic stress. These principles arise from cellular requirements to maintain electroneutrality, pH homeostasis, and a steady energy flow by coordinating glycolytic and ETC reactions. Qualitative connections have been drawn between expression data and ion homeostasis in the literature. For example, an inhibited respiration is regarded as an early response to an alkalinized cytoplasm due to a sudden efflux of proton that accompanies potassium intake upon osmotic upshift [28]. This initial response is followed by a rapid accumulation of glutamate—a potassium counterion—through glutamate importers or its biosynthetic pathways [26, 29]. Charge and pH neutrality are resumed in phase II upon flux regulations targeting the ETC, glycolysis, potassium and sodium transporters, glutamate transporters, glutamate biosynthesis, and glutamate decarboxylase. Here, the apparent inconsistency between the downregulated glutamate biosynthesis, observed from early stages [25], and elevated glutamate concentration is indicative of an active glutamate importer, such as glutamate/*γ*-aminobutyrate antiporter, in phase I.

To better understand the role of glutamate in osmoregulation, we computed the intrinsic proton- and charge-consumption (Eqs. (37) and (36); STAR Methods) of glutamate-biosynthesis pathways (*i.e*., GLUDy, GLUSy, and GLUDC) and the glutamate/*γ*-aminobutyrate antiporter. We found that glutamate accumulation through these biosynthetic pathways does not affect the net cytoplasmic charge. It also lowers the proton concentration of an already alkalinized cytoplasm in phase I. In contrast, glutamate uptake through the glutamate/*γ*-aminobutyrate antiporter results in an inflow of negative charge, which can counterbalance the accumulated K^+^ in the cytoplasm during phase I without affecting the pH, demonstrating the osmoregulatory role of glutamate antiporters (see Glutamate Role in Osmoregulation for a more detailed discussion).

#### Graphical representation

The hydrogen (Fig. 3C) and magnesium (Fig. 3D) maps graphically visualize equilibrium shifts with respect to pH and pMg perturbations. Similarly to the charge map, these maps can help elucidate regulatory responses to pH- and pMg-related stress conditions. Consider the hydrogen map for example. Here, a change in pH_*c*_ affects equilibrium constants, reaction Gibbs energies, and reaction rates according to La Chatelier’s principle [22]. If a reaction produces protons (Δ_*r*_*N*_H,*j*_ *<* 0) (Fig. 3C, blue arrows), then a positive perturbation in pH increases the equilibrium constant 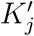, which, in turn, shifts the reaction more in the forward direction, so that protons are produced at a higher rate to counter the effect of the change. The magnesium map can be interpreted in a similar manner.

#### Recap

The ABCs shape transcriptional regulatory responses to environmental stresses. The combined effects of ion-binding and charge-balance constraints strongly restrict possible regulatory strategies to restore electroneutrality and pH homeostasis upon osmotic shock. In particular, our analysis revealed why regulating the cytoplasmic charge through glutamate antiporters is more desirable than glutamate-biosynthesis pathways. This highlights the importance of the GAD operons in osmoregulation, the transcriptomic function of which has been recently detailed using large datasets and machine learning [30].

### 2.5. Proton and charge imbalances underlie transcriptional regulatory responses to acid stress

#### pH *homeostasis is a fundamental cellular process*

Bacteria can survive extreme acidic environments thanks to several sophisticated acid-resistance mechanisms, such as glucose-repressed-oxidative, glutamate-dependent, and arginine-dependent systems [31, 32]. Acid stress causes increased proton entry into the cell that disrupts charge and the proton balances. The cytoplasmic pH is perturbed as a result, affecting equilibrium constants, reaction energies, and reaction rates. This invokes transcriptional regulatory responses, similar to those observed in osmoregulation discussed above. Being directly involved in pH-homeostasis and reaction equilibria, reaction proton-consumption discussed in the previous section are naturally expected to play a key role in acid-stress regulation.

#### Transcriptional responses to pH *changes*

To better understand how pH-dependent equilibrium shifts can affect transcriptional responses, we sought to identify connections between reaction proton-consumptions and gene expressions in phase-I/II RNA-sequencing data [32]. We identified 41 differentially expressed genes (*>* 1.4-fold change within an hour after pH down-shift) with the same regulatory targets (except phosphate transporters) in the reduced network as osmotic stress. However, there are key differences in regulatory responses between the two conditions: Contrary to osmotic stress, NADH dehydrogenase is upregulated, while phosphate and osmoprotectant transporters are downregulated under acid stress. These gene expression changes are consistent with the regulatory objective to neutralize the acidified cytoplasm by adjusting proton inflow and outflow rates.

For any given reaction, we found the intrinsic proton-consumption to be a reasonable indicator of its transcriptional regulatory state. Interestingly, among the 14 reactions with the highest proton consumption or production (|Δ_*r*_*N*_H_| *>* 1 mol-H/mol-rxn) in the hydrogen map (Fig. 3C), 11 are associated with the differentially expressed genes in phase I [33] and II [32]. To quantify the extent to which reaction proton-consumption in the reduced network can influence acid-stress response, we generated a connectivity map to identify associations between key metabolic pathways and transcriptional regulators (Fig. 4). Glycolysis, glutamate metabolism, and the ETC are major regulatory targets in the reduced network. Their extrinsic proton consumption (Δ_*r*_*N*_H_, units mol-H/gDW/h) in the base state are -21.23 (glycolysis), 7.18 (glutamate biosynthesis), and 113.78 (ETC). This analysis yielded three important findings.

First, Glycolysis is another major regulatory target. Overall, it is upregulated under acid and osmotic stresses. Interestingly, it comprises reactions with high intrinsic proton production (GAPD and PFK) and consumption (PYK). The same is true of charge consumption and production. As a result, the net proton consumption of the entire pathway (Δ_*r*_*N*_H_ = −21.23 mol-H/gDW/h) is much smaller than that of the ETC (Δ_*r*_*N*_H_ = 113.78 mol-H/gDW/h), even though the fluxes (obtained from FBA analysis) passing through these pathways are comparable. This feature allows the cell to reverse the flux direction through glycolysis in response to other simultaneous stresses (*e.g*., carbon-source change) without a significant disruption to pH homeostasis.

Second, the glutamate-dependent system has been reported as the most effective acid-resistance mechanisms, complementing primary responses [32]. Glutamate metabolism contains some of the highest proton consuming reactions in the reduced network, a possible explanation for their observed upregulation. They also carry high flux (especially GLUDy) whether or not the cell is under stress [28], so they can effectively attenuate acidification of the cytoplasm. The upper TCA cycle is downregulated, further promoting this mechanism by directing the flux of precursors from glycolysis and lower TCA towards glutamate biosynthesis.

As previously stated, these pathways are also involved in osmoregulation. However, unlike in pH regulation, GLUDy and GLUSy are both downregulated in response to osmotic stress. This raises a question as to why GLUDC is upregulated during osmoregulation when pH ≳7.5. Here, contrasting transcriptional responses to osmotic and acid stresses in relation to the proton balance might offer a plausible answer: Under osmotic stress, K^+^ is the main glutamate counterion when the cytoplasm is alkalinized in phase I with GLUDC counterbalancing proton influx through sodium antiporters and osmoprotestant symporters in phase II. In contrast, under acid stress, the proton becomes one of the main glutamate counterions when the cytoplasm is acidic in phase I with GLUDC, GLUDy, and GLUSy offsetting proton leakage through the membrane in phase II.

Third, the ETC can contribute the most to proton balance in the cell. However, it is directly or indirectly coupled to many crucial cellular processes such as membrane potential and redox balance. Therefore, regulation strategies that solely target the ETC would partially disrupt essential cellular functions and reduce the robust flexibility of bacteria to adapt to extreme and diverse environmental perturbations.

Thus, application of the ABCs allows for the biophysical interpretation of transcriptional regulatory responses to stimuli, such as acid stress.

### 2.6. Thermodynamic constraints drive the evolution of high-affinity phosphate transporters

#### Different ABCs have different evolutionary implications

By examining individual ABCs in isolation, one can identify the most dominant constraints under a given condition. Such identification may have important evolutionary implications because each of these constraints could have been dominant at different point in time during evolution. For example, thermodynamic laws are believed to have constrained the evolution of biochemical-reaction networks ever since the formation of primitive cells to ensure the energy requirements and spontaneity of prebiotic chemical reactions [34]. However, the osmotic-pressure differential and membrane potential likely have changed significantly during the course of evolution from ion-permeable porous membranes in protocells to ion-impermeable lipid-bilayers in modern cells [35]. This forms a rational basis for chronological postdictions concerning the evolution of alternative pathways.

#### Low- and high-concentration phosphate transporters

*E. coli* has two phosphate-transport systems (Fig. 5E), namely *Pit* (low-affinity) and *Pst* (high-affinity). Here, we sought to determine whether any of the ABCs could have been restrictive enough to hinder the operation of the *Pit* system in phosphate-limited environment, thereby, driving the evolution of the *Pst* system. These transporters, which differ in their stoichiometry (Fig. 5E), are of particular interest because inorganic phosphate is an important constituent of energy carrying molecules (*e.g*., ATP), cofactors (*e.g*., NADH), and information-storage molecules (*e.g*., DNA). Therefore, *E. coli* maintains an optimal homeostatic phosphate level around 1–10 mM [37, 38].

#### Is is possible to achieve cytoplasmic phosphate concentrations above 1 mM while satisfying all ABCs?

We computed feasible concentration ranges for the reduced network, in which either the *Pit* or *Pst* system was the only active phosphate transporter. We compared these feasible ranges at two periplasmic concentrations, representing phosphate-limited (⟦pi _p⟧ _1_) and phosphate-rich (⟦pi-p⟧_2_) environments (Fig. 5C). While the feasible concentration range for the cytoplasmic phosphate is almost insensitive to its periplasmic concentration when the *Pst* system is active, achieving cytoplasmic concentrations above 1 mM is impossible in phosphate-limited environments when the *Pit* system is active. Moreover, comparing the upper bound on the cytoplasmic phosphate concentration provided by the ABC and TMFA at the forgoing periplasmic concentrations reveals that the operation of the *Pit* system is limited by the thermodynamic constraints, not the osmotic-balance or other charge-related constraints (Fig. S6; STAR Methods).

#### High-affinity transporters maintain sufficient phosphate influx in phosphate-limited environment to support growth

We compared the transport kinetics of the two phosphate transport systems (Fig. 5D) in a wild-type strain of *E. coli* to demonstrate the transition between the *Pit* and *Pst* systems in response to phosphate-starvation stress. Clearly, *E. coli* can remain functional in a wide range of extracellular concentrations, importing phosphate through the *Pit* system at high rates when it is available in excess. However, when the

*Pit* system reaches its equilibrium at low extracellular concentrations, *E. coli* exhibits a stress response by taking up phosphate through the *Pst* system to retain the phosphate influx, albeit at lower rates. Note that, this is not the case for every transmembrane protein in *E. coli*. Cytochrome oxidase in the ETC is an example, where the transition from low-affinity (cytochrome-bo3) to high-affinity (cytochrome-bd) system occurs most likely due to kinetic reasons. Because these reactions are both highly energetic in their forward direction (Δ_*r*_*G*^′º^ ≈ − 90 kJ/mol), their equilibria are never reached for any biologically feasible concentration ranges of ubiquinone, ubiquinol, and oxygen. Thus, the thermodynamic constraints are unlikely to be responsible for the low activity of cytochrome-bo3 in oxygen-limited environments.

#### Recap

The ABCs can explain the principles underlying the evolution of multiple transport systems. Specifically, we showed that the thermodynamic constraints restrict the operation of the *Pit* system in phosphate-limited environments, necessitating the evolution of higher affinity transport systems.

## 3. Discussion

> Those [constraints] that operate at any given level are still valid at all more complex levels — Francois Jacob [39]

Biological functions are subject to myriad constraints. The constraints on the function of abiotic and early biotic cells have been the subject of much discussion from Darwin’s warm-pond hypothesis to the role of geothermal vents at the bottom of the ocean. Fundamental to this discussion is the role of abiotic constraints. Given Francois Jacobs’ quote, these constrains are fundamental and affect the function of ancestral and modern cells. The ABCs, thus, apply to functions across the biosphere.

In this study we (i) formulated the ABCs, (ii) developed a computational platform to apply them, (iii) elucidated regulatory targets that shape transcriptional regulatory responses to external stresses, (iv) showed that measured quantitative metabolomes satisfy these constrains, (v) found that they predict thermodynamic bottlenecks that mirror isotope labelling experiments, and (vi) elucidated the reasons underlying the evolution of multiple phosphate transporters. These results show the broad applications of the ABC to understand biological functions and their origins. In fact, the two are not separable.

Constraint-based approaches have significantly advanced in recent years, incorporating more detailed molecular descriptions of biological networks [4]. Despite their expanded predictive scope, these approaches still fall short in one key regard: Metabolite concentrations do not enter the formulation through governing equations or constraints grounded in first principles. To fill this gap, we introduced a constraint-based approach to characterize the concentration state of metabolic networks. This approach predicts feasible concentration and Gibbs-energy ranges resulting from the constraints biological networks are subject to.

Besides characterizing the metabolome, abiotic constraints shape regulatory responses to various stress conditions. We studied osmotic- and acid-stress regulations as illustrative examples to establish associations between transcriptional responses and reaction proton- and charge-consumptions. We argued that the net proton and charge consumption of the electron transport chain, glutamate biosynthesis, and glutamate transporters can effectively counteract the proton and charge imbalances induced by osmotic and acid stresses, providing an explanation for why they are subject to transcriptional regulation. Other reasons for why certain reactions are regulatory targets have been proposed. For example, reactions with a large thermodynamic driving force are believed to be under stricter transcriptional regulatory control than near-equilibrium ones since their rates are less sensitive to concentration perturbations [10]. Reaction rates that are sensitive to variations in concentrations can be effectively controlled through allosteric regulation. Hence, near-equilibrium reactions are expected to rely less on transcriptional regulations than highly exergonic ones.

An observed stress response is generally a result of overlapping allosteric and transcriptional regulatory processes, the organization and mechanism of which depend on: (i) regulation objective (*e.g*., restoring homeostasis after a vital constraint is violated), (ii) regulation constraints (*e.g*., charge and H^+^ balance, osmotic balance, cell structural integrity, gene clusters and physical proximity, ATP and redox balance), and (iii) regulatory targets (biochemical reactions). We showed several ways, in which flux adjustments to biochemical reactions can modulate critical intracellular variables (net charge, proton and magnesium-ion concentrations, metabolite concentrations) to alleviate stress-induced imbalances, providing a more comprehensive description of stress-specific regulatory responses. The gene-expression activity of glycolytic reactions in response to acid stress is an example, where *pfkB* and *pykA* are differentially expressed. Although the associated reactions have the largest thermodynamic driving force, they also produce and consume the highest amount of H^+^ among glycolytic reactions. Therefore, their transcriptional regulation is possibly more related to their role in pH homeostasis than their energetics.

We showed that the ABCs are consistent with metabolomic data across carbon sources. To investigate whether intracellular concentrations are associated with optimality principles, we studied two cellular objectives that have been previously considered [18], namely thermodynamic and enzyme-saturation efficiency. We found that biochemical reactions do not generally satisfy these objectives. Specifically, we identified several reactions from the TCA cycle and glycolysis that operate near their equilibrium during the exponential growth phase. Near-equilibrium reactions lower dissipative energy loss at higher protein cost. They also offer functional advantages, enabling rapid flux-direction reversal in energy-metabolism pathways to robustly achieve new steady states upon a wide variety of environmental perturbations. These observations underscore the complex and multifaceted nature of cellular metabolism, which cannot be fully described by single-objective or oversimplified optimality principles [40].

We studied the role of abiotic constraints in the evolution of phosphate transport systems. These concepts may be applied in a broader context to prebiotic reaction networks to gain insights into the origins and evolution of early metabolism [41, 34]. The fundamental constraints we implemented in our analysis are particularly relevant to origins-of-life theories. Specifically, the membrane potential, transmembrane ion gradients, charge balance, and thermodynamic laws are believed to have constrained the most ancient pathways for the formation of small organic molecules [34]. Several scenarios have been suggested for the origins of primitive life [42]. For all these theories to be plausible, one must establish a balance between the availability of an energy source that could have existed on the early earth (*e.g*., proton gradient in hydrothermal vents) and the energy required to operate the most energy demanding step (*e.g*., methyl synthesis) of the proposed chemistry for carbon metabolism (*e.g*., CO_2_ reduction by H_2_) [34]. These hypotheses can be rigorously formulated and examined within the quantitative framework we developed in this work to address some of the fundamental questions concerning the emergence of life from prebiotic chemistry.

The detailed formulation of the ABCs unraveled additional layers of attributes associated with biochemical reactions besides energetics and stoichiometry that are important to network functions. These constraints can quantify perturbations in charge, pH, and pMg homeostasis induced by biochemical reactions, explaining the commonalities and differences in regulatory processes with similar objectives. Electroneutrality and pH homeostasis are examples of basic regulatory objectives that underlie transcriptional regulatory response to acid and osmotic stress. We elucidated the associations between these stress responses and charge-related constraints. Furthermore, the ABCs exhibit consistency with a variety of metabolomic, isotope-labeling, and differential expression data, laying the foundation for integrated models of flux, concentration, and macromolecular expression in the future that are capable of predicting functional states under any growth or stress conditions.

Taken together, the ABC shows that biological interactions with the inorganic world are foundational to the evolution and function of cells, and, thus, should be accounted for in the studies of fundamental biological processes.

## Author Contributions

### Acknowledgment

We would like to thank Sharon Grubner and Jonathan Hsu. We also appreciate Jared Broddrick’s valuable comments on the manuscript. This work was funded by the Novo Nordisk Foundation (Grant Number NNF10CC1016517), the National Institutes of Health (Grant Number R01GM057089), and the Institute for Systems Biology’s Translational Research Fellows Program (J.T.Y.).

## STAR *\** Methods

### Reactants and Species in Biochemical Reactions

Most metabolites behave as weak acids in the cell, donating proton to the intracellular fluid in multiple steps [22, Chapter 1]. The resulting protonation states can, in turn, bind to several metal ions (*e.g*., K^+^ and Mg^2+^) in separate steps. These charge states, referred to as *species*, can coexist in the cell with varying distributions depending on the pH and ionic strength of the solution. Several species can be biologically active under physiological conditions, simultaneously participating in biochemical reactions. In general, all the protonation states and their magnesium-bound counterparts are active in the cell, especially, those with phosphate groups [43, Section 9.4.2]. We account for all the active charge states of a given metabolite, incorporating proton-dissociation and magnesium-binding reactions (up to two Mg^2+^)

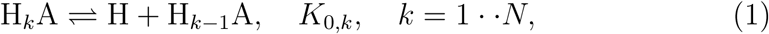

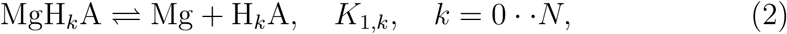

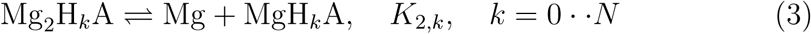

into our formulation. Here, *N* is the number of proton dissociation steps and A represents the minimum charge state of a metabolite in the cell with *K*_0,*k*_, *K*_1,*k*_, and *K*_2,*k*_ the respective proton-dissociation and magnesium-binding constants. These reactions tend to run faster than enzymatic reaction, always remaining close to their equilibrium [22, Chapter 1]. This allows to determine the distribution of species independently from the extension of biochemical reactions. Therefore, we simplify our analysis by lumping all the species together, representing them as a single *reactant*, with respect to which the abiotic constraints are to be expressed

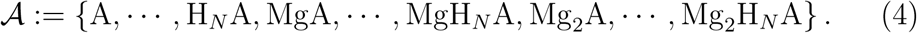

**Figure S1:**
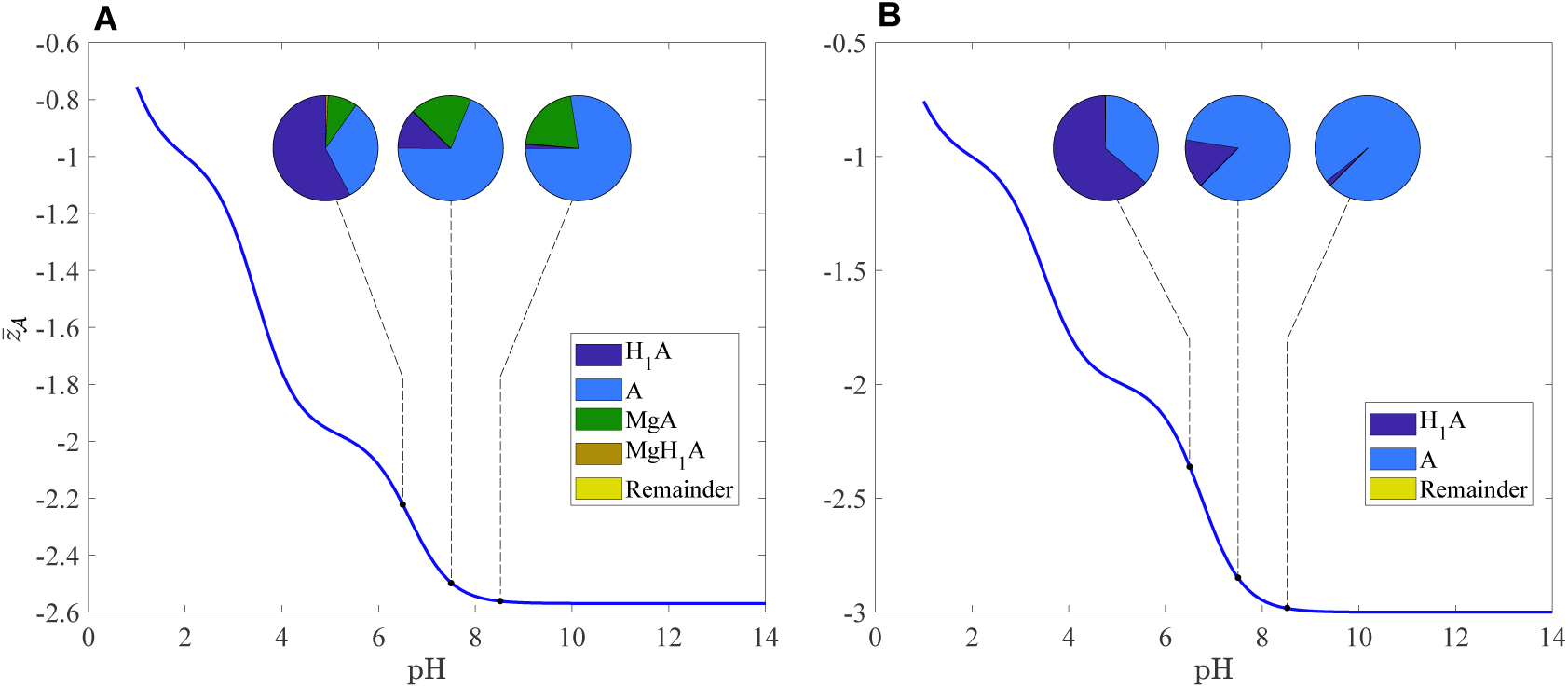
Effective charge of D-glycerate 2-phosphate, related to Fig. 1. (A) *I* = 300 mM and pMg = 2.699. (B) *I* = 300 mM and pMg = 16. Pie charts indicate the distribution of dominant species at the respective pH.

Here,*𝒜* represent a reactant in the cell, the concentration of which is given by

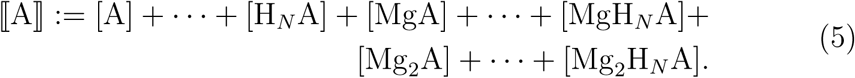

For each species A^′^ ∈ *𝒜*, the corresponding model fraction *ρ*_A_*′* := [A^′^]*/* ⟦A⟧ is calculated from proton-dissociation and magnesium-binding constants, irrespective the reaction A^′^ participates in (see Composition of Reactants). Accordingly, we define an effective charge

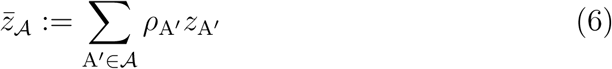

for the reactant *𝒜*. Here, *z*_A_*′* denotes the charge of the species A^′^. This definition ensures the consistency of formulations with respect to reactants and species by requiring 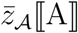 to furnish the total charge of all the species in *𝒜*. Although species always have an integral charge, reactants can generally carry fractional charges depending on the ionic strength, pH, and pMg of the solution because of their dependence on the mole fractions (Fig. S1). The effective buffer intensity 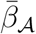, effective charge squared 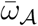, effective proton content 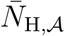, and effective magnesium content 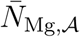 of *𝒜* are defined similarly

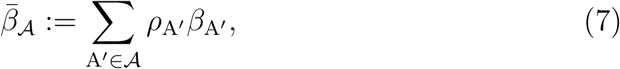

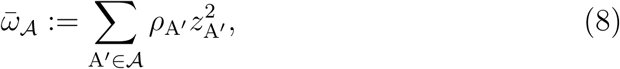

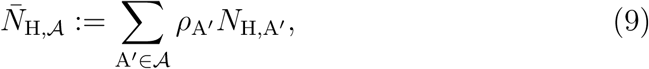

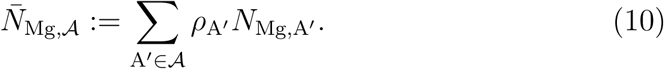

In this article, the terms *metabolite* and *reactant* are interchangeably used. To distinguish between individual charge states and their aggregate representation in our formulation when necessary, we specifically refer to a metabolite and its charge states as reactant and species, respectively.

**Figure S2:**
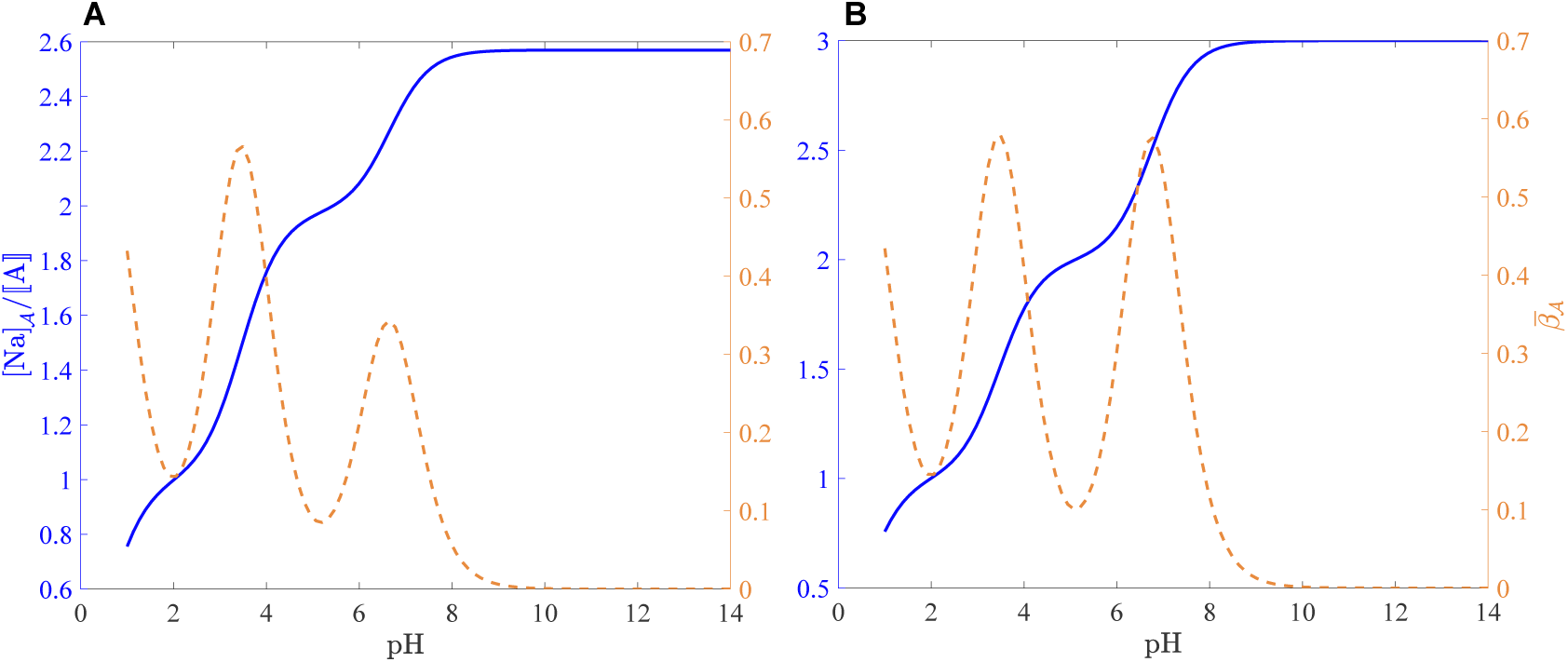
Strong-base equivalent (solid) and buffer intensity (dashed) of D-glycerate 2-phosphate, Related to Fig. 1. (A) *I* = 300 mM and pMg = 2.699. (B) *I* = 300 mM and pMg = 16.

### Buffer Capacity and Buffer Intensity

The buffer capacity *ℬ* is a measure of how much a strong base (or acid), such as NaOH, is needed to increase the pH of a solution. The buffer capacity of a mixture of reactants can be obtained from a superposition of the buffer capacities of individual reactants [44]. Therefore, we first derive an expression for the buffer capacity of a reactant *𝒜*. We consider all the charge states of *𝒜* arising from proton-dissociation and magnesium-binding reactions Eqs. (1)-(3) in our derivation. The charge balance for this system reads

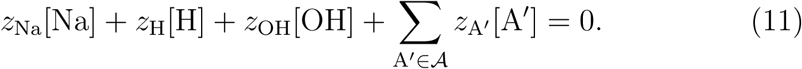

Because NaOH is a strong base, it completely hydrolyzes, so that [Na] represents the concentration NaOH needed to neutralize *𝒜*. Note that, we did not explicitly state the charge that each ion carries in the this equation for brevity—a notational convention we adopt throughout this document. We decompose [Na] into a water [Na]_*w*_ and reactant component [Na] _𝒜_, which we refer to as the strong-base equivalent of *𝒜*, according to [Na] = [Na]_*w*_ +[Na] _𝒜_, where

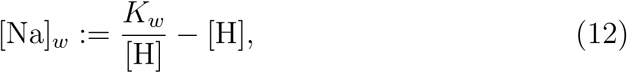

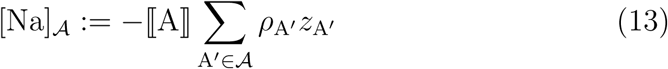

with *K*_*w*_ the water dissociation constant. Accordingly, the buffer capacity of water and reactant *𝒜* are defined [44]

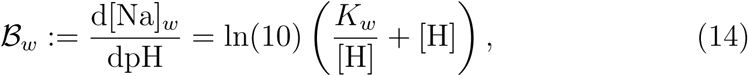

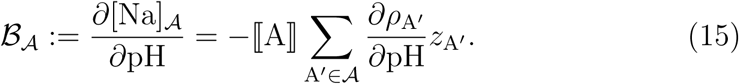

The effective buffer intensity of *𝒜* is simply defined as 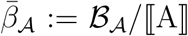. The buffer capacity can be approximated by linearizing *ℬ*_A_ around a pH of interest. It is correspondingly defined as the amont of a strong base one should add to a solution to increase its pH by one. Figure S2 illustrates how the strong-base equivalent and buffer intensity of *𝒜* vary with pH. The total buffer capacity of a mixture of reactants is determined from those of the individual reactants

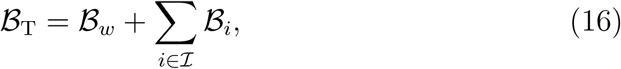

where *ℐ* is the index set of reactants in the solution.

### Osmotic Pressure and Activity Models

Osmosis is a prevalent phenomenon in electrolyte systems, where there are ion-concentration differentials. Consider the osmotic coefficient of a multicomponent aqueous solution [45]

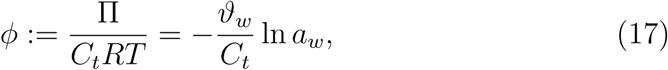

where *C*_*t*_ is the total molar concentration of solutes, *R* gas universal constant, and *T* temperature. Here, *ϑ*_*w*_, *a*_*w*_, and *x*_*w*_ denote molar density, activity, and mole fraction of water. The osmotic pressure Π measures the pressure of the solution relative to that of pure water at the same temperature. Using Pitzer’s model for the water activity [46]

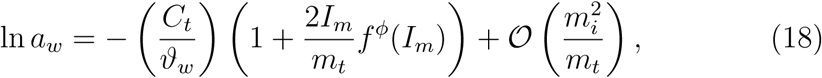

where

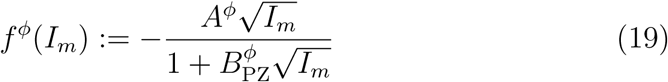

with *A*^*φ*^ = 0.391475 kg^1*/*2^ mol^−1*/*2^, 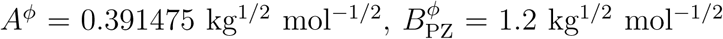, *m*_*t*_ total molal concentration of solutes, and *I*_*m*_ the molal ionic strength [46]. We neglect the second and higher order terms in *m*_*i*_, corresponding to ion-ion interactions, in Eq. (18). Since the parameters required to estimate these interactions are not generally known for biological systems, they are usually neglected [22]. Satisfactory results have been reported in equilibrium studies of biochemical reactions using activity models based on this approximation in concentration ranges of physiological relevance, justifying this assumption [22]. Molal concentrations can be converted to molar concentrations according to 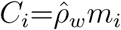, where *C*_*i*_ and *m*_*i*_ are molar and molal concentrations of solute *i*. Moreover, 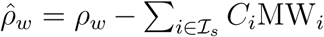, where 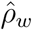 and *ρ*_*w*_ are the reduced density and density of solution with *ℐ*_*s*_ and MW_*i*_ the index set of solutes in the solution and molecular weight of solute *i*. In general, these densities are functions of temperature, pressure, and solute concentrations. However, in biological systems, they are commonly approximated by the density of water since solute concentrations in the cell are negligible compared to water [22]. We adopt this approximation to convert between molar and molal concentrations through a multiplicative constant. Note that, if a molar ionic strength *I* is passed to *f* ^*φ*^, the constants in Eq. (19) must be adjusted according to 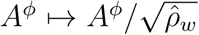 and 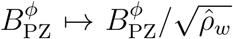. Given the relationship 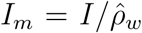, these ensure the consistency of ionic-strength and parameter units. Substituting Eq. (18) in Eq. (17), we derive an expression for the osmotic coefficient

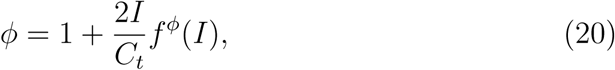

where molality-to-molarity conversion has been applied. We use this expression to estimate the osmotic pressure of the cytoplasmic and periplasmic fluids in our model

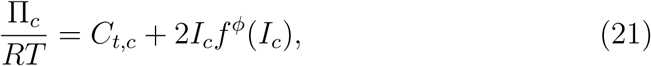

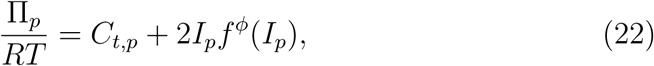

where the subscripts *c* and *p* denote the quantities associated with cytoplasm and periplasm.

The activity coefficients of solutes are also needed for equilibrium computations that will be discussed in subsequent sections. The activity coefficient of charge solutes in ionic solutions can be generally expressed as

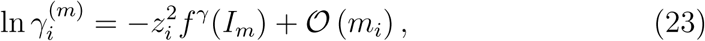

where 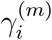 is the molality-based activity coefficient. Two widely accepted activity models are the extended Debye-Hückel and Pitzer, respectively given by [22, 46]

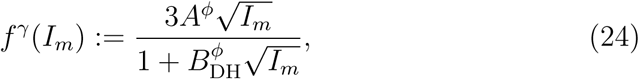

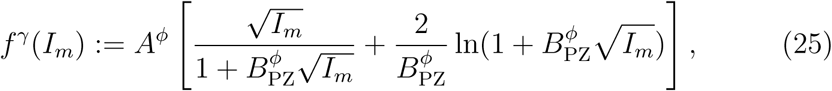

where 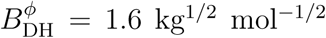 [47]. As with the osmotic coefficient, the parameter transformations 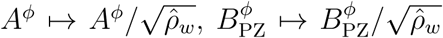, and 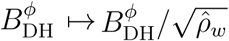 apply if *I* is passed to *f* ^*γ*^ instead of *I*_*m*_. Our formulations in the forthcoming sections are with respect to molarity-based activity coefficients *γ*_*i*_. These are related to molality-based activity coefficients by 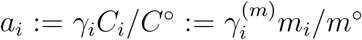, where *C*º = 1 mol/L-w and *m*º = 1 mol/kg-w are the standard concentrations. As will be discussed, activity coefficients are needed to calculate the formation energies of species at a given *I* from standard formation energies, which we approximate using group-contribution methods [48]. These techniques estimate group-contribution parameters using the extended Debye-Hückel model by minimizing the difference between the predicted and measured equilibrium constants of biochemical reactions. Since equilibrium constants are highly sensitive to these parameters, we adhere to the extended Debye-Hückel model in our formulation.

### Abiotic Constraints

Metabolism is a hallmark of living systems. It is orchestrated by a delicate balance between metabolite concentrations, biochemical-reaction fluxes, and macromolecular levels in the cell. Metabolic networks are tightly intertwined with complex, overlapping regulatory and signaling networks, enabling the cell to sustain homeostasis or adapt to new environments by transitioning between homeostatic states [12]. Quantitative description of dynamical systems of such complexity using mechanistic kinetic models is often restricted to small networks [49] due to computational challenges, limited kinetic data, and incomplete knowledge about the mechanisms of enzymatic reactions [50].

We introduce a constrained-based formalism to characterize the metabolome. Rather than specifying a unique time-dependent concentration state, we seek a set of concentration states that respect all the constraints biological networks are subject to, allowing for a unified treatment of steady-state and oscillatory dynamics. Fundamental constraints, such as mass-action principle, charge balance, and osmotic balance, imposed by the laws of thermodynamics, electroneutrality, and osmotic pressure, are obeyed by all electrochemical systems. The mass-action principle arises from the dissipative structure of chemical-reaction systems [51, 52], while the charge and osmotic balances are essential for cell-volume and ion-transport stability [53]. Evolutionary constraints, such as fixed ionic strength and buffering capacity, have emerged from the evolutionary adaptation of biological networks. The ionic strength plays a crucial role in several cellular processes, such as osmoregulation, enzyme activity, and protein structure [54], while the buffering capacity is associated with pH homeostasis [55]. These condition-specific parameters are controlled by regulatory networks to maintain a stable metabolism under various charge-related stress conditions.

The complexities of regulatory networks impede the application of mechanistic dynamic models, even for those controlling the simplest cellular processes [12]. Therefore, we adopt a top-down approach, where the phenotypes arising from important regulatory processes (*e.g*., osmoregulation and ion homeostasis) are implicitly accounted for by incorporating them as additional constraints (the evolutionary constraints) into our model. The forgoing fundamental and evolutionary constraints are mathematically expressed as

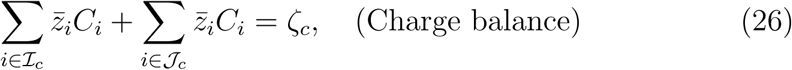

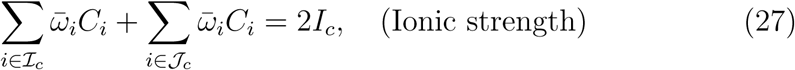

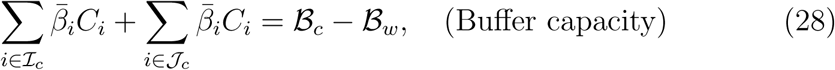

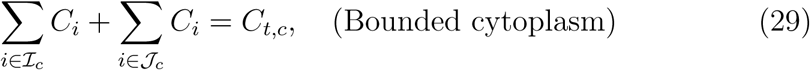

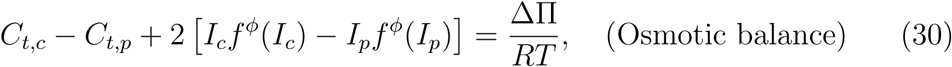

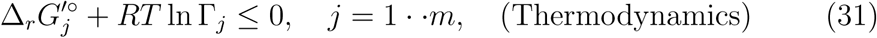

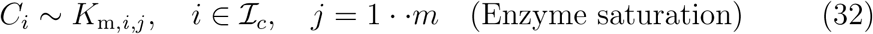

with *C* the vector of cytoplasmic reactant concentrations (*C*_𝒜_ and ⟦A⟧ are interchangeably used to denote the concentration of *𝒜*. For a reactant with only one charge state, [A] and ⟦A⟧ are indistinguishable), *C*_*t,c*_ total cytoplasmic concentration, *C*_*t,p*_ total periplasmic concentration, *ζ*_*c*_ net cytoplasmic charge, *I*_*c*_ cytoplasmic ionic strength, *I*_*p*_ periplasmic ionic strength, *ℬ*_*c*_ total cytoplasmic buffering capacity, *ℬ*_*w*_ buffering capacity of water, ΔΠ := Π_*c*_ − Π_*p*_ osmotic-pressure differential, *f* ^*φ*^ Pitzer’s function for osmotic coefficient [46, Eq. (5)], Δ_*r*_*G*^′º^ standard transformed Gibbs energy of reaction, Γ reaction quotient, *K*_m_ Michaelis constant, *j* reaction index, *i* reactant index, and *m* number of reactions. Moreover, *ℐ*_*c*_ and *𝒥*_*c*_ are the index sets of cytoplasmic reactants and transmembrane ions (K^+^, Na^+^, Cl^−^, Mg^2+^, OH^−^, and H^+^) [56] affecting the membrane potential Δ*ψ*. Similarly, the index sets of periplasmic reactants and transmembrane ions are denoted *ℐ*_*p*_ and *𝒥*_*p*_. We applied these constraints to characterize the CSS of a reduced metabolic network of *E. coli* (Fig. S3).

**Figure S3:**
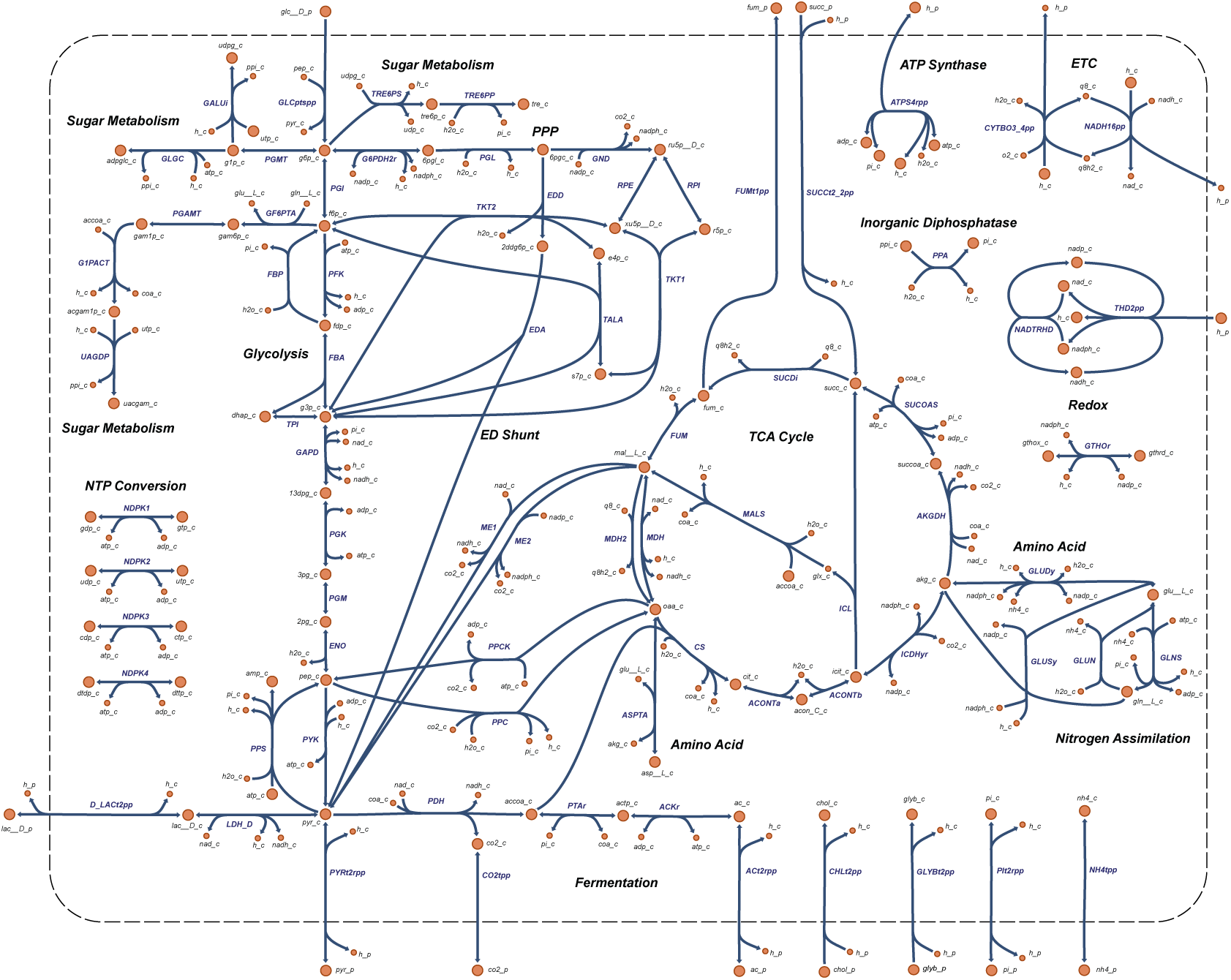
Metabolism in *E. coli*. A reduced metabolic network, comprising more than 90% of the entire metabolome by mole, is constructed to simulate growth on four carbon sources, including glucose, acetate, pyruvate, and succinate.

**Figure S4:**
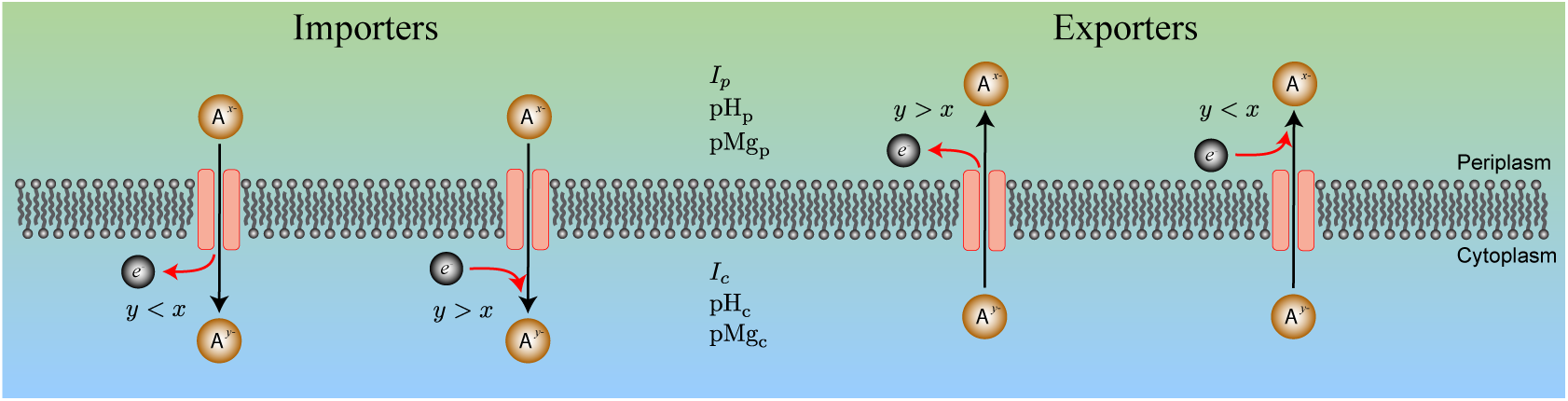
Convention adopted to calculate the reaction charge-consumption of transporters, Related to Fig. 1. Charge exchange between reactants and water is assumed to take place on the product side of transport reactions.

Equations (26)-(31) represent charge balance, fixed ionic strength, fixed buffering capacity, bounded cytoplasmic metabolome, osmotic balance, and mass-action principle, respectively. Equation (32) corresponds to the enzyme-saturation constraint (see Enzyme Kinetics), which is written as a scaling expression here because it does not generally hold for every reaction in the network (*e.g*., amino-acid degradation reactions [11]). The standard transformed energy 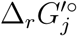 of reaction *j* is obtained from the standard transformed formation energy 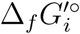 of reactants *i* ∈ *ℐ*_*c*_ ∪ *ℐ*_*p*_ according to their coefficients in the stoichiometric matrix *S*. The intrinsic reaction proton-consumption Δ_*r*_*N*_H,*j*_, magnesium-consumption Δ_*r*_*N*_Mg,*j*_, and charge-consumption Δ_*r*_*Z*_*j*_ are similarly defined. Reactions with Δ_*r*_*Z*_*j*_ *>* 0 and Δ_*r*_*Z*_*j*_ *<* 0 consume and produce negative charge in the cytoplasm by convention, respectively. In the main text, we generally refer to Δ_*r*_*Z*_*j*_ as the reaction charge-consumption, irrespective of its sign. When necessary, we specifically refer to it as charge consumption or charge production to emphasize the sign of Δ_*r*_*Z*_*j*_. The same conventions apply to Δ_*r*_*N*_H,*j*_ and Δ_*r*_*N*_Mg,*j*_. We use the same notation to denote extrinsic reaction proton-, magnesium-, and charge-consumption. These are defined as the product of their intrinsic counterparts and the respective flux. We clarify whichever is relevant in the context by explicitly stating their units (intrinsic and extrinsic consumptions are measured in mol/mol-rxn and mol/gDW/h, receptively).

Since biochemical reactions involve transformations between reactants with multiple charge states that exchange ions with the cytoplasmic and periplasmic fluids, they generally do not balance charge, proton, and magnesium ions. Indeed, the reaction charge-, proton-, and magnesium-consumption discussed above are defined to quantify this property. In the following discussion, we make these definitions more precise. Starting with charge consumption, we introduce three components for Δ_*r*_*Z*_*j*_

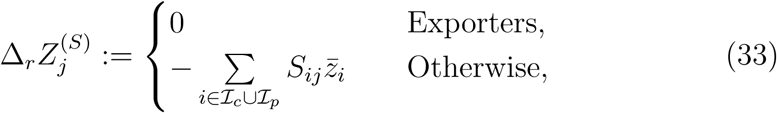

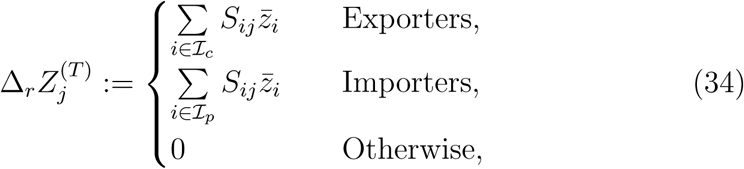

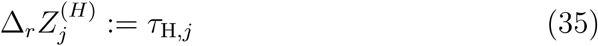

with *τ*_H,*j*_ the number of translocated protons in reaction *j*, where *τ*_H,*j*_ = 0 for intracellular reactions, *τ*_H,*j*_ *>* 0 if proton is translocated from the periplasm to cytoplasm, and *τ*_H,*j*_ *<* 0 if proton is translocated from the cytoplasm to periplasm. Exporters are thermodynamically spontaneous in the direction, where reactants are transferred from the cytoplasm to periplasm, and importers are thermodynamically spontaneous in the opposite direction. Note that, 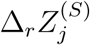 reflects the amount of charge exchanges between reactants and water due to proton dissociation and magnesium binding. For intracellular reactions, charge exchanges occur within the cytoplasmic fluid. However, ambiguity may arise for transport reactions as to whether reactants exchange charge with the cytoplasmic or periplasmic fluids. To avoid this ambiguity, we assume that these exchanges occur on the product side of transport reactions (Fig. S4). The definition given by Eq. (34) is in accordance with this assumption. The reaction charge-consumption is defined

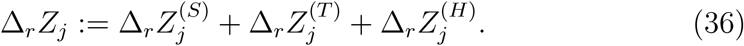

We term 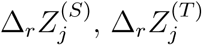, and 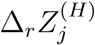 the stoichiometry, reactant-transport, and proton-transport components. According to this definition, Δ_*r*_*Z*_*j*_ does not affect the overall charge balance of the cytoplasmic fluid for intracellular reactions. However, for transport reactions, it measures the net negative charge flowing into or out of the cell.

Reaction proton-consumption is an important quantity, elucidating the role of biochemical reactions in pH homeostasis. To properly defined it, a key difference between Δ_*r*_*Z*_*j*_ and Δ_*r*_*N*_H,*j*_ must be accounted for: While the goal of Δ_*r*_*Z*_*j*_ is to capture how biochemical reactions affect reactant-cytoplasmic fluid charge transfer and the overall charge balance of the cell, Δ_*r*_*N*_H,*j*_ is defined to measure the contribution of a reaction to pH_*c*_. This is based on the assumption that maintaining a near-neutral pH_*c*_ is a more essential constraint for the operation of biological networks than the overall proton balance of the cell. Accordingly, reaction proton-consumption is defined by two components

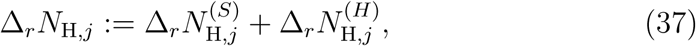

where

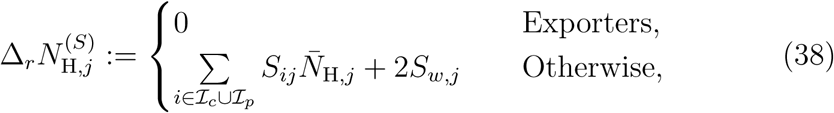

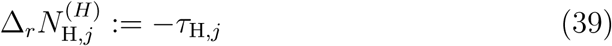

with *S*_*w,j*_ the stoichiometric coefficient of water in reaction *j* (see Transformed Reaction Gibbs Energy). As with charge consumption, proton exchanges between reactants and water is assumed to occur occur on the product side of transport reactions in Eq. (38). Reaction magnesium-consumption is defined in a similar manner

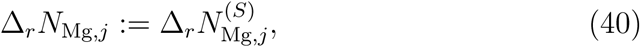

where

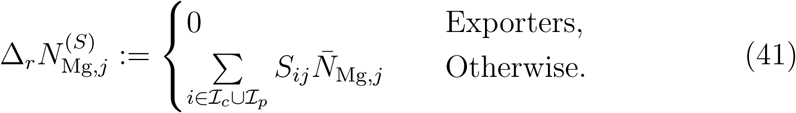

In Eqs. (26)-(32), the concentration of cytoplasmic reactants, excluding the transmembrane ions, are the unknowns. The transmembrane-ion concentrations, *I*_*c*_, and *ℬ*_*c*_ are condition- and strain-specific, the values of which are specified at the outset (Table S2). All other quantities are the given parameters of the model (Table S2). Periplasmic concentrations are assumed to be identical to those of the growth medium with a specified composition (M9 minimal medium [57], detailed in Table S1).

We define the CSS as a subset of the concentration space, where all the biophysical constraint are satisfied:

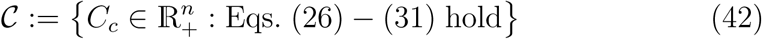

with *n* := |*ℐ*_*c*_|, *n*^′^ := |*ℐ*_*p*_|, and *C*_*c*_ a subvector of *C* corresponding to *ℐ*_*c*_. Geometrically, the CSS can be represented as the intersection of the affine subspace defined by the equality constraints Eqs. (26)-(30) and the thermodynamically concentration solution space corresponding to Eq. (31) (Fig. 1B). We characterize the CSS, which can generally be nonconvex and disconnected, using global optimization and sampling techniques. The first furnishes global bounds on metabolite concentrations and reaction energies, regardless of whether the CSS is connect or not. The second provides expectations of concentrations *𝔼*(*C*_*i*_) and reaction energies 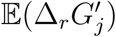, standard deviations of concentrations *σ*(*C*_*i*_) and reaction energies 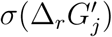, and violation probabilities of thermodynamic constraints *P*_*j*_. Here, we first identify an interior point *q* by solving a parametric optimization problem (Eq. (85)) that maximally distances *q* from the thermodynamic constraints, simultaneously avoiding arbitrarily small concentrations, which are biologically irrelevant. We found these solutions to be well-correlated with metabolomic data reported in the literature. We then explore a neighborhood of *q* by generating curves along random directions inside the CSS to sample the space and determine the thermodynamic constraints that are violated more frequently (see Characterization of Concentration Solution Space).

We reformulate Eqs. (26)-(31) with respect to mole fractions to arrive at a compact dimensionless form

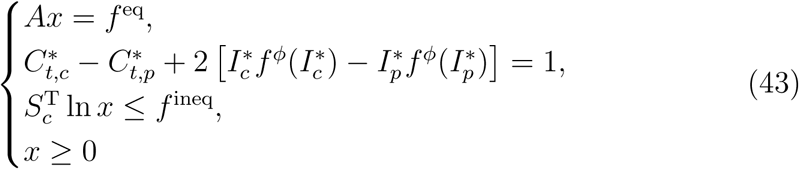

with

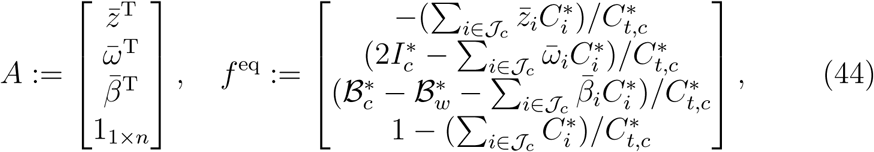

and

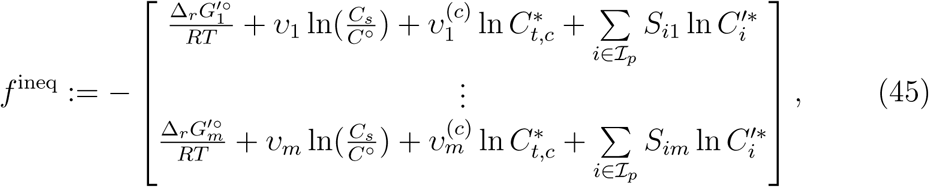

where all the concentrations, ionic strengths, and buffer capacities are scaled with *C*_*s*_ := ΔΠ*/RT*. Here, *x* := *C*_*c*_*/C*_*t,c*_ is the vector of cytoplasmic mole fractions, the superscript ^*^ denotes scaled parameters or scaled variable, *C*^′^ is the vector of periplasmic reactant concentrations, *S*_*c*_ is a submatrix of *S* containing all the rows corresponding to 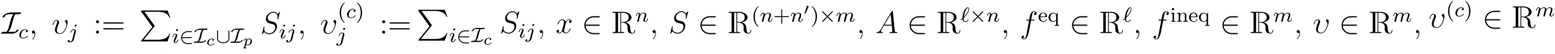 and *S*_*c*_ ∈ ℝ^*n*×*m*^ with *l* the number of linear equalities (*i.e*., rows of *A*) in Eq. (43). Note that, because *I*^*^ is passed to *f* ^*ϕ*^ instead of *I*_*m*_ in Eq. (43), the constants of Eq. (19) must be adjusted according to 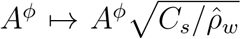 and 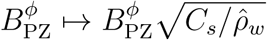. In Eq. (43), *x* and 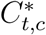 are unknown variables; all the other parameters are known and specified at the outset. The CSS in the mole-fraction space is defined

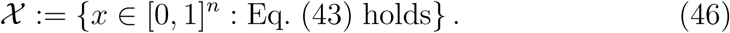

Since the CSS spans several orders of magnitude in the concentration and mole-fraction space, we use the logarithmic map Ξ := *x ↦ y* = ln *x* to reformulate Eqs. (43) and (46) into the forms

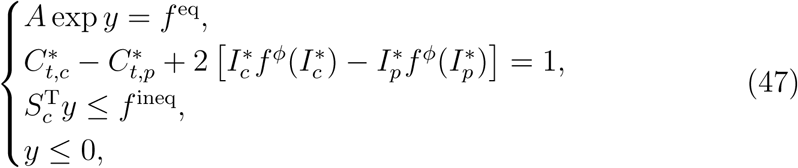

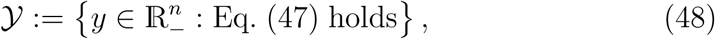

in which it is more computationally tractable to characterize the CSS. We refer to the mole-fraction and logarithmic mole-fraction spaces as the *X* and *Y* space, respectively.

### Enzyme Kinetics

The reversible Michaelis–Menten mechanism [58]

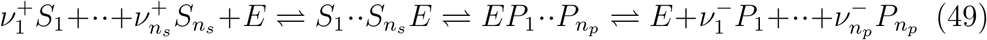

is one of the simplest mechanisms proposed to study the kinetics of biochemical reactions. It leads to the separable rate law [58]

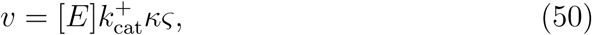

where

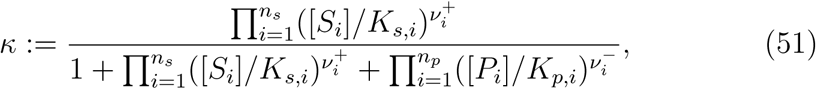

and

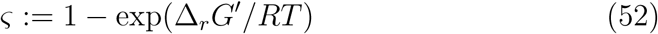

are the enzyme-saturation and thermodynamic efficiencies. Here, *K*_*s,i*_ and *K*_*p,i*_ are the Michaelis constants of the substrates and products with [*E*] the total enzyme concentration, 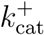 the turnover number, *n*_*s*_ the number of substrates, and *n*_*p*_ the number of products. It is clear from Eq. (50) that to draw maximum flux at minimum protein cost, both efficiencies must be maximized. However, the maximum saturation efficiency (*i.e*., *κ* → 1 achieved when [*S*_*i*_] ≫ *K*_*s,i*_ and [*P*_*i*_] ≫ *K*_*p,i*_) cannot be realized by every reaction in the network because an unbounded increase in all substrate and product concentrations may conflict with other constraints imposed on the network, such as osmotic balance and molecular crowding. Therefore, a more relaxed efficiency criterion is usually used, where a reaction is considered efficient when it is at least half-saturated in one direction ([*S*_*i*_] ≥ *K*_*s,i*_ in the forward or [*P*_*i*_] ≥ *K*_*p,i*_ in the backward direction) [11, 18]. Reversible reactions are required to be half-saturated or more in both directions to be considered efficient. For reactant *i*, we define the maximum Michaelis constant

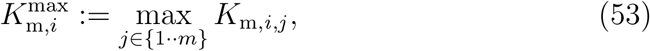

where *K*_m,*i,j*_ stands for *K*_*s,i*_ and *K*_*p,i*_ in Eq. (51) that are associated with reaction *j*. If the reactant *i* does not participate in the reaction *j*, then *K*_m,*i,j*_ = 0. Accordingly, the enzyme-saturation-efficiency criterion for the entire network is defined

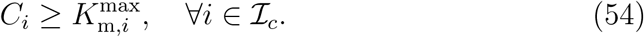

If Eq. (54) is not satisfied for reactant *i*, it implies that at least one of the reactions it participates in is undersaturated.

### Composition of Reactants

We discussed in previous sections how to represent the abiotic constraints with respect to effective quantities. These were defined as weighted averages of the respective quantity for species constituting a reactant. The mole fractions of species were the weights in these definitions. Here, we show how to compute these mole fractions from binding constants for a reactant *𝒜*. First, we derive expressions to relate proton dissociation and magnesium binding constants at a given ionic strength to those evaluated at standard condition (infinite dilution at the same pressure and temperature, where *I* → 0)

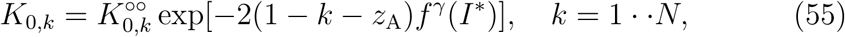

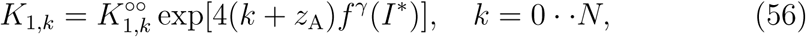

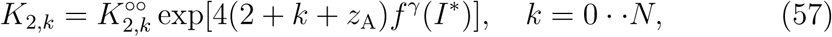

where 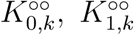, and 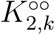 are reference equilibrium constants, and *z*_A_ is the charge of the minimum charge state of *𝒜* (see Reactants and Species in Biochemical Reactions). Moreover, we define *K*_0,0_ := 1 for notational convenience. Next, we introduce the binding polynomial

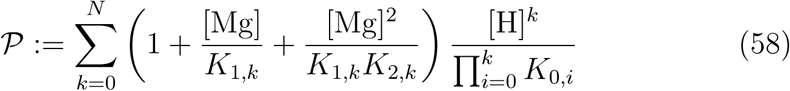

for proton-dissociation and magnesium-binding equilibria given by Eqs. (1)-(3). Finally, we obtain the species mole fractions

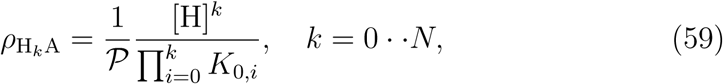

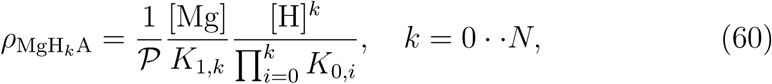

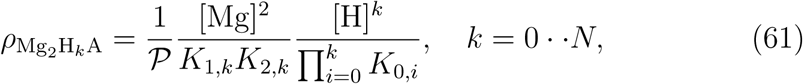

### Formation Energy of Species from Formation Energy of Minimum Charge State

Before proceeding to calculate the formation energy of reactants, we provide useful expressions to ascertain the formation energy of species from that of the minimum charge state. The formation energies of all the charge states arising from Eqs. (1)-(3) can be obtained from 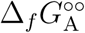 according to

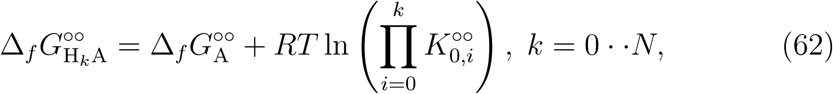

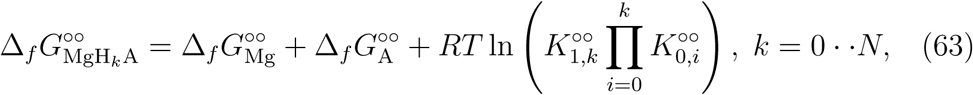

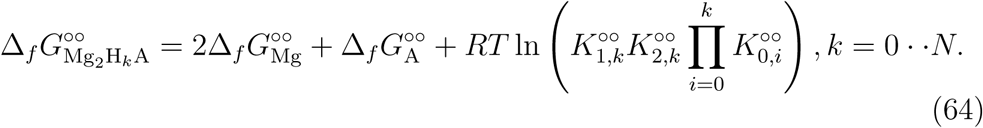

Note that, in deriving these equations, we substituted the reference formation energy of proton 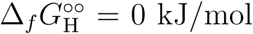, and explicitly expressed the reference formation energy of magnesium ion 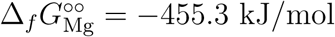 [22].

### Transformed Formation Energy of Reactants

Following the development of Alberty [22], the transformed Gibbs energy of formation of the species A^t^ ∈ *𝒜* is written

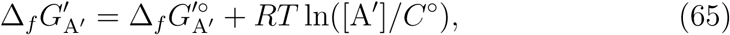

where 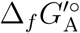 is the standard transformed Gibbs energy of formation evaluated at the same ionic strength, pH, and pMg, accounting for all the nonidealities and external force fields. It can, in turn, be expressed as

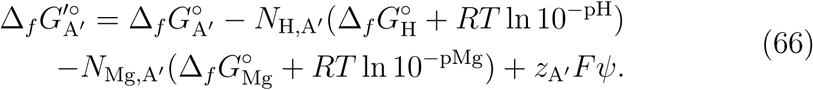

The individual components of 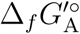 are given by

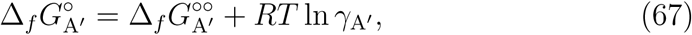

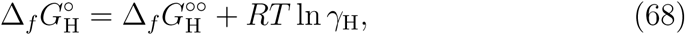

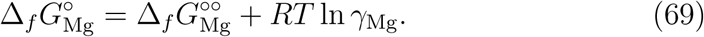

Here, *F* is the Faraday constant, is the external potential field at the point where Gibbs energy is evaluated, and the superscript ^ºº^ denotes the reference state of Gibbs energy at infinite dilution of the respective species. The standard formation energy of reactant *𝒜* is expressed with respect to that of species

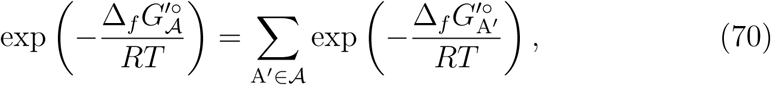

which can be reformulated to

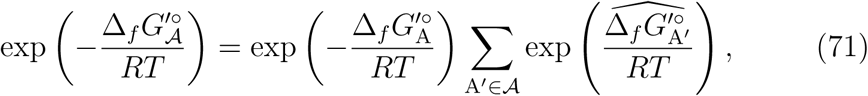

where 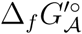 is the standard formation energy of *𝒜*, 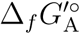 the standard formation energy of the minimum charge state, and 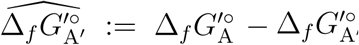. Substituting Eqs. (66)-(69) in Eq. (71), we arrive at

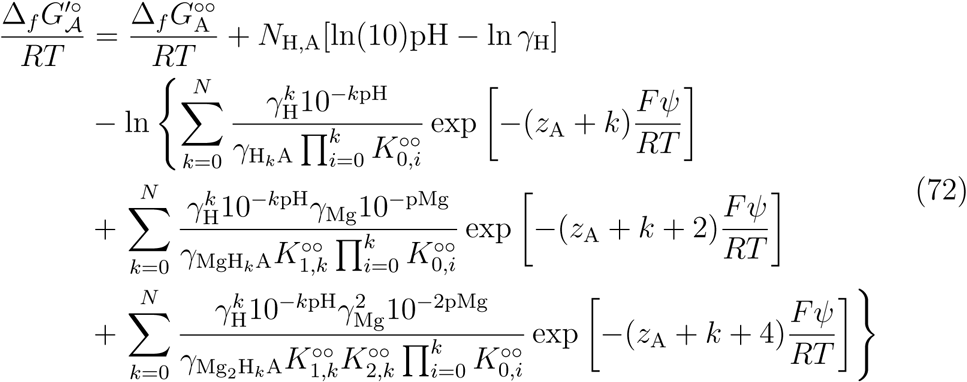

with *N*_H,A_ and 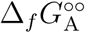 denoting the proton content and reference formation energy of the minimum charge state. Note that, the second and third sums in Eq. (72) correspond to magnesium-bound states, while the first sum arises from protonation states. We estimated the the transformed Gibbs energy of reactions with and without magnesium-bound states. However, all the results presented in the main text were computed by neglecting the charge states arising from magnesium bindings. As previously stated, we obtained reaction energies using the reference formation energies of species furnished by group-contribution methods [48]. These methods tune their parameters based on expressions that only account for protonation states. Therefore, using these formation energies in Eq. (72) with all the charge states included introduces errors into reaction-energy computations, which are no longer bounded by the guaranteed uncertainty bounds of group-contribution methods.

### Transformed Reaction Gibbs Energy

The standard transformed reaction Gibbs energies and apparent equilibrium constants are obtained from the transformed formation energies of reactants

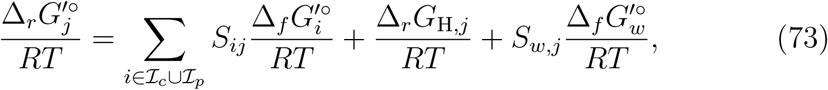

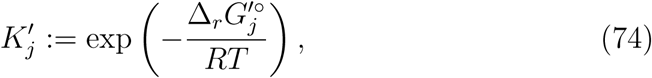

where *j* is the reaction index, and

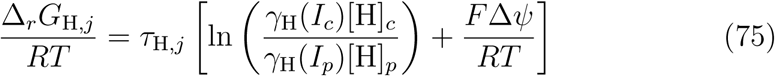

is the proton motive force associated with reaction *j* with Δ*ψ* := *ψ*_*c*_ − *ψ*_*p*_ the membrane potential. The transformed Gibbs energy of reaction *j* is

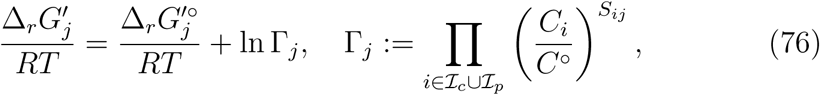

where Γ_*j*_ is the quotient of reaction *j*. Note that, in this formulation, the concentrations of proton and water do not contribute to the reaction quotients nor do their stoichiometric coefficient to *S*. Instead, the concentrations and stoichiometric coefficients of water and proton are implicitly accounted for in Eq. (73). The standard transformed formation energy of water is given by

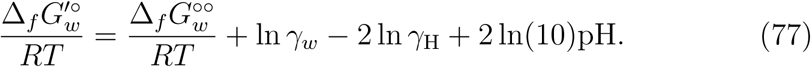

In biological systems, *γ*_*w*_ ≈ 1 because the concentration of reactants are negligible compared to water [22]. In group-contribution methods, *γ*_*w*_ is assumed to be a constant and lumped into the reference formation energy 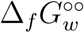, and the resulting expression is used to fit the model parameters (*i.e*., formation energies of species at infinite dilution) to equilibrium data. In this work, we adopt 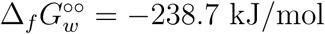 [48].

### Growth Medium and Model Parameters

As previously stated, the concentration of cytoplasmic metabolites, excluding the transmembrane ions, are the only unknowns in Eqs. (26)-(32). The goal in our model is to characterize the CSS for any given growth medium, the composition and parameters of which are fully specified at the outset. The results presented in the main text all describe the metabolic state of *E. coli* growing in an M9 minimal medium (Table S1) during the exponential growth phase. From the composition of the growth medium, we computed the concentrations of all the periplasmic ions, ascertaining effective charges and *I*_*p*_.

**Table S1:** Composition of M9 minimal medium, Related to Fig. 1.

| Compound | Concentration | Dilution factor |
| --- | --- | --- |
| Na<sub>2</sub>HPO<sub>4</sub> | 33.9 g/L | 0.2 |
| KH<sub>2</sub>PO<sub>4</sub> | 15.0 g/L | 0.2 |
| NH<sub>4</sub>Cl | 5.0 g/L | 0.2 |
| NaCl | 2.5 g/L | 0.2 |
| CaCl<sub>2</sub> | 0.0001 M | 1 |
| MgSO<sub>4</sub> | 0.002 M | 1 |
| CO<sub>2</sub> | 0.0001 M | 1 |
| Choline | 0.0001 M | 1 |
| Acetate | 0.0001 M | 1 |
| Glycine betaine | 0.001 M | 1 |
| Carbon source | 5 mg/L | 1 |

Besides the composition of the growth medium, several other parameters must be specified at the outset to determine intracellular concentration ranges using Eqs. (26)-(32). We chose characteristic parameters associated with the exponential growth of *E. coli* (Table S2) to study feasible intracellular concentrations, and how they are restricted by abiotic constraints.

### Characterization of Concentration Solution Space

In this section, we present two methods for characterizing the CSS constructed by the abiotic constraints outlined previously. First, we compute global bounds on cytoplasmic concentrations and reaction energies using global optimization techniques to determined their respective biologically feasible ranges. The lower bounds are obtained by solving

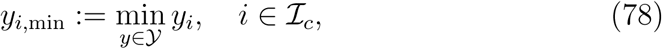

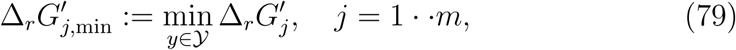

where *𝒴* is given by Eqs. (47) and (48). The upper bounds are similarly determined. Note that, in the concentration space, the feasible concentration ranges can span several orders of magnitude possibly over a disconnected CSS. Therefore, it is more convenient to perform global optimization computations in the *Y* space. Second, we compute expectations and standard deviations of concentrations and reaction energies. Given a random variable *X* : *𝒞* ↦ ℝ defined over the CSS, the expectation and standard deviation of *X* are defined as

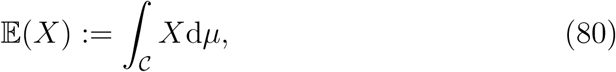

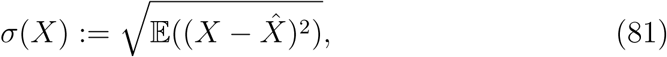

where 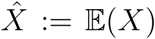, and *µ* is a probability measure of *X*, encapsulating all the intracellular dynamics and the related parameter uncertainties (*e.g*., kinetic parameters, protein structure, enzyme saturation, and spacial gradients). This probability measure, which is expected to be condition specific, is poorly understood in biological systems. Without any information about the probability measure for the organism of interest, the most natural choice for *µ* is the volume measure. However, *𝒞* as previously stated, is generally a nonconvex and disconnected set, rendering volume integrals difficult to compute. Therefore, we further simplify our analysis by using a line measure to approximate expectations and standard deviations. We compute these line integrals along random curves generated in the equality-constraint manifold (Fig. S5A and B). Since all these computations are performed in the *Y* space, we first describe briefly the mathematical formulation of the CSS in this space.

**Table S2:** Model parameters. Intracellular concentrations of transmembrane ions are approximated using values reported in the literature [59, 60]. These parameters characterize the growth of *E. coli* during the exponential phase, Related to Fig. 1.

| Parameter | Value | Unit | Parameter | Value | Unit |
| --- | --- | --- | --- | --- | --- |
| $T$ | 25 | $^{\circ}\text{C}$ | $I_c$ | 300 | mM |
| $\Delta\Pi$ | 303 | kPa | $\mathcal{B}_c$ | 62.5 | mM |
| $\Delta\psi$ | -140 | mV | $\zeta_c$ | 0 | mol-$e$/L |
| $\hat{\rho}_w$ | 1 | kg/L | $[\text{K}]_c$ | 200 | mM |
| $\text{pH}_p$ | 7.0 | — | $[\text{Na}]_c$ | 14 | mM |
| $\text{pH}_c$ | 7.5 | — | $[\text{Cl}]_c$ | 20 | mM |
| $V_{\text{cell}}$ | 2.6 | fL | $[\text{Mg}]_c$ | 2 | mM |

Let *ℳ* denote the abstract (*n* − *l*)-dimensional manifold in the *Y* space defined by the linear equalities in Eq. (43) and 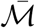 its isometric embedding in ℝ^*n*^ furnished by the embedding map *ι* : *ℳ* → *Y* (Fig. S5C). We chose this particular embedding to simplify the construction and computation of the foregoing line measures. Moreover, let *𝒜* := *A*^T^, *b* := (*f* ^eq^)^T^, and

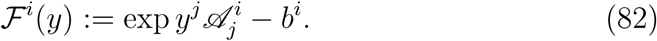

Note that, in previous sections, we did not distinguish between subscripts and superscripts to denote the components of vectors and covectors. Vectors were treated as *n*-tuples, the components of which were represented by subscripts (column vector in matrix form) without reference to the space they were associated with. However, in this section, the components of vectors and covector are denoted by superscripts and subscripts, which are represented by row and column vectors in matrix form, respectively. The same convention applies in general to covariant and contravariant components of tensors. We also adopt the Einstein summation rule, where repeating a dummy index as a superscript and subscript implies summation over an appropriate range of the index. Assuming that *A* has full rank (*i.e*., rank(*A*) = *l* with *l < n*), we have

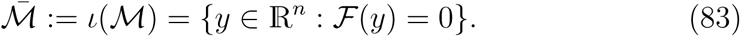

The tangent space at a point *y* on 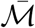 is given by 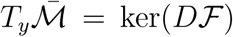 [61]. In matrix form, the derivative of *ℱ* can be written *D ℱ* = *AE*, where *E* := diag(exp *y*). Explicit expressions can be derived to construct coordinate charts *ϕ* for *ℳ* using null-space bases of *A*. Let *N* ∈ ℝ^*n*×(*n*−*l*)^ be a matrix containing an orthonormal basis of ker(*A*). Then, Φ : ↦ *χ* ln(*x* + *χN* ^T^) with *χ* ∈ ℝ^*n*−*l*^ contains information about the coordinate chart *ϕ* at the point *y* = ln *x* because Φ = *ι* º *φ*^−1^. The components of *χ* can be regarded as the coordinates of *ϕ* at the point *y*. We can now define an induced metric [62] for *ℳ* according to

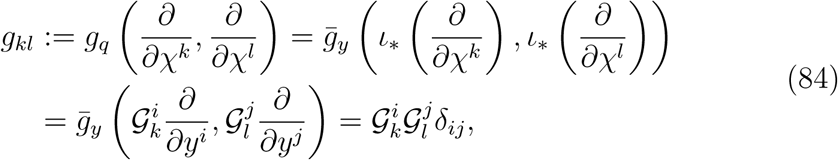

where *δ* is the Kronecker delta tensor, *y* = *ι*(*q*), and 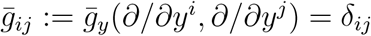 is the standard Euclidean metric that *Y* ≡ ℝ^*n*^ is equipped with. Here, *ι*_*_ is the derivative of *ι*, furnishing 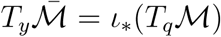, and 𝒢 := *D*Φ^T^ = *N* ^T^*E*^−1^ in matrix form.

**Figure S5:**
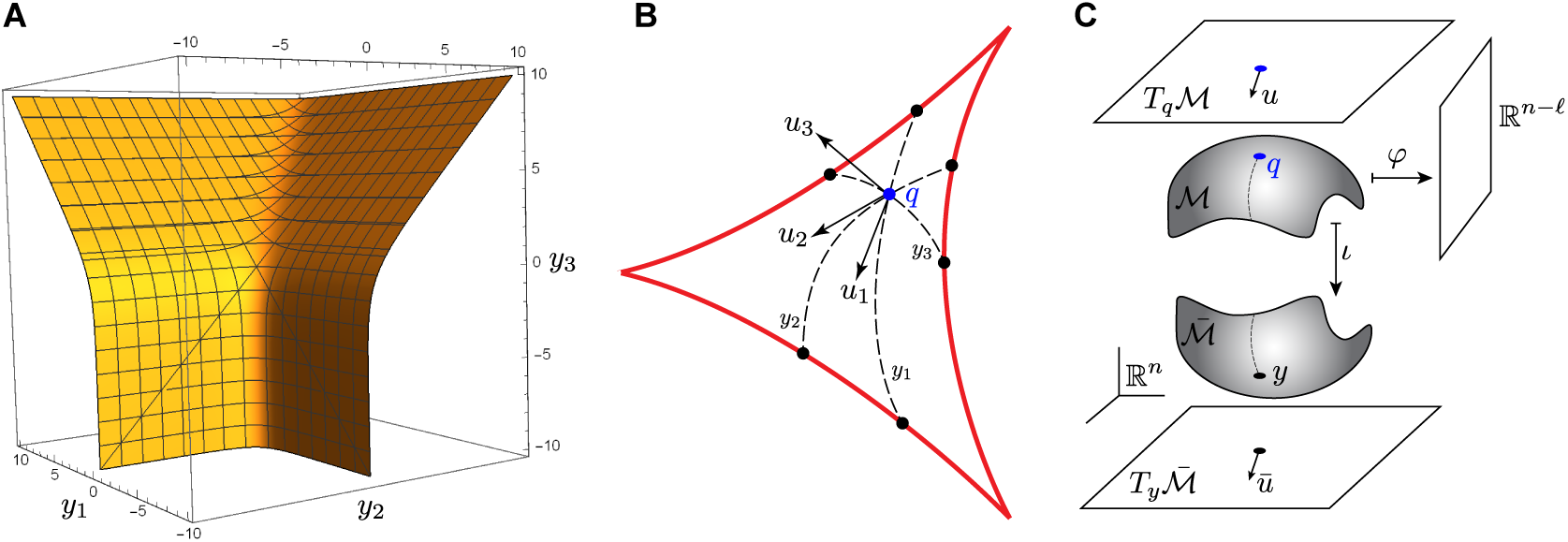
Geometry of the CSS in the *Y* space, Related to Fig. 2. (A) A two-dimensional example of equality-constraint manifold 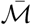 embedded in ℝ^3^, where *A* = [1 1 − 1] and *f* ^eq^ = 1. (B) Schematic representation of the trajectory-tracing method used to characterize the CSS. Trajectories are constructed in 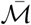 from an interior point *y* along vectors *ū*_1_, *ū*_2_, and *ū*_3_ that are randomly generated in the tangent space 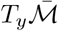 and continued until they cross at least one of the thermodynamic constraints (red curves). The expectation and standard deviation of any function defined on *𝒞* are ascertained by computing the respective line integrals along these trajectories. (C) Schematic representation of the equality-constraint manifold *ℳ*, tangent space at *q*, and their embeddings in ℝ^*n*^.

Before generating random curves in 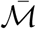, we first require a feasible point of the CSS, preferably far away from all its boundaries, in the *Y* space. We generate a one-parameter family of interior points by solving

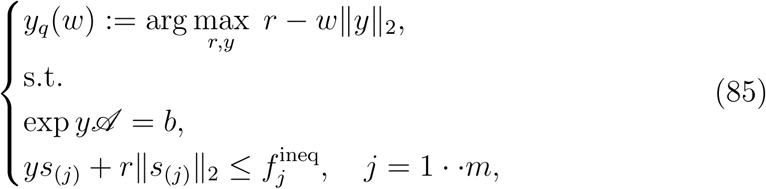

where *s*_(*j*)_ denotes the *j*th column of *S*_*c*_. Note that, to derive Eq. (85), we solved the second equation in the system Eq. (47) for 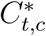, the value of which is substituted in *f* ^eq^ and *f* ^ineq^, so it is not explicitly included in the system Eq. (85). Observe that, *y*_*q*_(0) is the Chebyshev center [63, Section 8.5.1] of the polyhedron defined by the thermodynamic constraints in the *Y* space that is restricted to 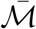. For *w >* 0, the objective function identifies an interior point of this polyhedron that is maximally distanced from all the thermodynamic constraints with respect to the second norm in the *Y* space, while avoiding arbitrarily large negative values for *y*. This provides biologically relevant points inside the CSS.

Several approaches can be taken to generate random curves in 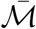 from *y*. Here, we outline a computationally tractable technique. First, a vector *u* = *u*^*k*^*∂/∂χ*^*k*^ is generated in *T*_*q*_ℳ using the basis *∂/∂χ*^*k*^ associated with the coordinate chart *ϕ* discussed before, where the components *u*^*k*^ ∈ [−1, 1] are random numbers. The embedding of *u* in 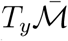 can be readily obtained according to *ū* = *ι*_*_(*u*) = *ū*^*i*^*∂/∂y*^*i*^, where 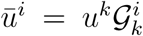. Next, a line 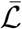 is constructed in 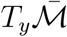 that passes through *y* along the tangent vector *ū* with the parametric representation 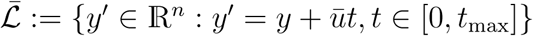. Finally, a trajectory 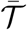 is constructed in 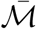 by finding the orthogonal projection of 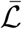 onto 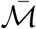. This is accomplished by solving

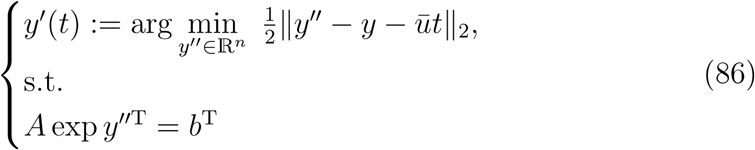

with

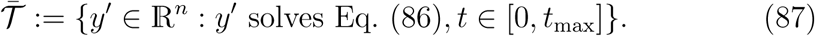

The Karush–Kuhn–Tucker (KKT) conditions [63, Section 5.5.3] for Eq. (86) reads

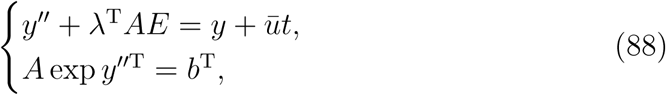

where *λ* is the dual variable corresponding to the equalities in Eq. (86). It is also a column vector associated with the co-normal space 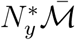. Moreover, *b* is associated with the normal space 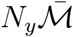 and is a row vector. All the other vectors are associated with 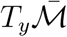 and are row vectors. Differentiating the KKT system Eq. (88) with respect to *t*, we arrive at

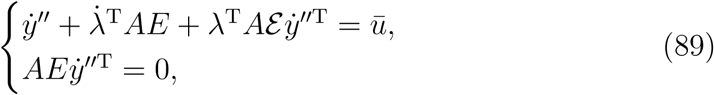

where overdot denotes derivatives with respect to *t*, and *ℰ* ∈ ℝ^*n*×*n*×*n*^ is defined as

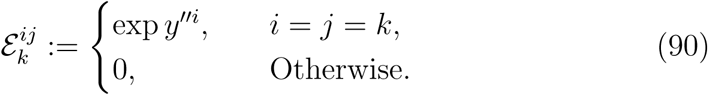

Equation (89) is integrated with respect to *t* subject to the initial conditions *y*″(0) = *y* and *λ*(0) = 0 until one of the thermodynamic constraints is violated. Note that, the constraint *y* ≤ 0 need not be checked directly because one of the equality constraints in Eq. (89) forces the sum of the metabolite mole fractions to be strictly less than one. As a result, *y*″^*i*^ = 0 can never be realized for any metabolites along trajectories constructed by solving Eq. (89). We formulated the global optimization problems Eq. (78), (79), and (85) in the General Algebraic Modeling System [64] and solved using the global solver BARON [65] equipped with the local nonlinear programming solver CONOPT and the linear programming solver CPLEX. Equation (89) was solved using standard time integrators in MATLAB R2019b.

Once random trajectories 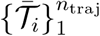 are constructed, the expectation of a random variable *X*, defined by Eq. (80), can be approximated

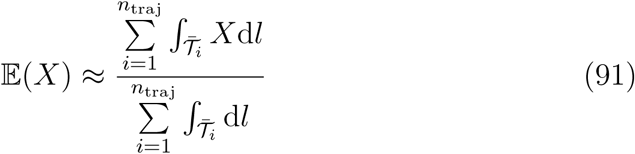

with the line measure 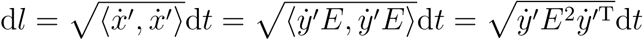. The standard deviation of *X* is similarly approximated. The violation probability of thermodynamic constraint associated with reaction *j* is given by

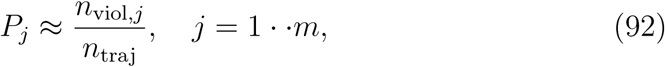

where *n*_viol,*j*_ is the number of trajectories intersecting the thermodynamic constraint of reaction *j*, and *n*_traj_ the total number of randomly generated trajectories. Expectations, standard deviations, and violation probabilities in Fig. 2 are estimated by generating 50000 trajectories in the CSS.

### Glutamate Role in Osmoregulation

To elucidate the glutamate-accumulation mechanism by which electroneutrality and pH homeostasis are achieved, we compared the intrinsic proton- and charge-consumption (see Eqs. (37) and (36)) of the impacted pathways. Interestingly, all major glutamate-biosynthesis pathways (*i.e*., GLUDy, GLUSy, and GLUDC) simultaneously consume proton and produce negative charge with similarly high intrinsic values (Δ_*r*_*N*_H_ ≈ 1 mol-H/mol-rxn and Δ_*r*_*Z* ≈ −1 mol-*e*/mol-rxn), whereas glutamate antiporters only produce negative charge (Δ_*r*_*N*_H_ ≈ 0 mol-H/mol-rxn and Δ_*r*_*Z* ≈−1 mol-*e*/mol-rxn). These features highlight the importance of glutamate antiporters during phase I when the cytoplasmic solution is alkalinized, allowing the cell to rapidly accumulate glutamate and potassium from the extracellular environment, so as to maintain electroneutrality without a significant disruption to pH homeostasis. Therefore, exporting excess glutamate, possibly through mechanosensitive channels (*e.g*., *mscL*) [26], when the cell is under no stress could be part of a robust regulatory network in bacteria, enabling them to create an extracellular glutamate pool, which can be used to flexibly handle various stress conditions causing ion imbalances.

Biochemical reactions can affect the cytoplasmic charge in two different ways, both of which are captured by our definition of Δ_*r*_*Z* (see Abiotic Constraints). Recall, Δ_*r*_*Z* reflects the amount of charge exchange between metabolites and the cytoplasmic fluid for intracellular reactions, while it measures the net charge transfer from periplasm to cytoplasm for transport reactions. If the overall cytoplasmic electroneutrality is regarded as a more essential osmoregulation objective than balanced intracellular charge exchanges, then it follows from our analysis that glutamate antiporters play a more significant role in glutamate accumulation during osmoregulation than glutamate-biosynthesis pathways—a conclusion supported by differential expression analyses of RNA-sequencing data.

### Flux Distribution

To compute extrinsic reaction charge-, proton-, and magnesium consumptions, the flux distribution across the network is required. We determine the flux state of the reduced network during the exponential growth phase using the genome-scale reconstruction model iML1515 [13]. To reliably estimate the metabolic fluxes, we identify the parsimonious solution of the resulting FBA model that best matches experimentally measured fluxes [12]. The parsimonious solution, furnished by minimizing a norm of the flux vector, postulates that the optimal flux state minimizes the total protein cost of the network, assuming that all biochemical reactions have the same protein cost. This objective has been shown to be associated with the flux state of several organisms under various conditions [18, 40], and, therefore, is adopted in our study. We formulate the FBA model as a bi-level optimization problem, incorporating the forgoing objectives. To simplify our analysis, we performed several numerical experiments and found that the upper-level objective, measuring the distance between the experimental and predicted fluxes, has the largest sensitivity to the upper bounds on the GLGC and NDPK1 fluxes in our reduced network. Accordingly, we determine the flux state by solving

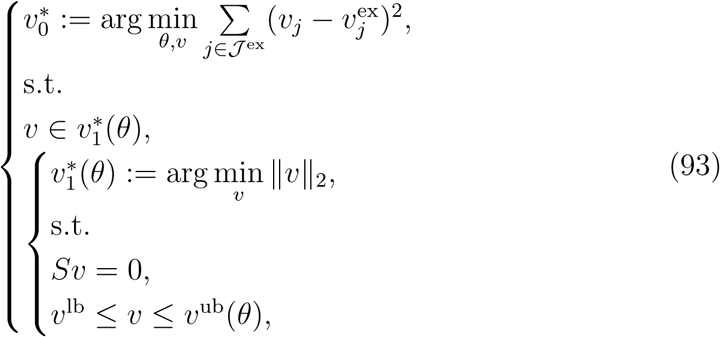

where *S* is the stoichiometric matrix of iML1515, *𝒥*^ex^ the index set of reactions with experimentally measured flux, *v*^lb^ and *v*^ub^ the lower and upper bounds on the flux vector, and *θ* a vector containing the upper bounds of GLGC and NDPK1. Note that, these upper bounds, which are decision variables in the upper level, are linearly expressed in the respective inequality constraints of the lower level. Thus, the lower-level problem of Eq. (93) is a convex quadratic programming problem, which we solve using the quadratic programing solvers of CPLEX. We solve the upper-level problem using gradient-descent methods.

### Feasible Concentration Ranges Affected by Phosphate Transport Systems

To determine the dominant constraints that restrict the operation of the *Pit* system at low periplasmic phosphate concentrations ⟦pi-p⟧, we compared the feasible concentration ranges of cytoplasmic phosphate predicted by BCS and TMFA at high and low periplasmic concentrations (Fig. S6). The upper bounds predicted by the BCS and TMFA models coincide at low and high periplasmic concentrations when *Pit* is active. In contrast, when *Pst* is active, the upper bound predicted by BCS is more restrictive than predicted by TMFA. These results indicate that the transition from the *Pit* to *Pst* systems with decreasing ⟦pi-p⟧ is induced by the *Pit* system becoming thermodynamically infeasible in phosphate-limited environments.

### Pathway Classification

In the main text, we classified reactants and reactions based on the pathway they participate in to contrast the energetics and metabolite distribution across major pathways in the reduced network. The abbreviations used to label these pathways in Figs. S3 and 2 are defined in Table S3.

**Table S3:** Pathway abbreviations, Related to Figs. S3 and 2.

| Abbreviation | Pathway |
| --- | --- |
| PPP | Pentose phosphate pathway |
| TCA | Tricarboxylic acid cycle |
| ED | Entner-Doudoroff Pathway |
| AA | Amino acid biosynthesis |
| Fer | Fermentation |
| NTP | Nucleoside triphosphate conversion |
| GB | Glycine betaine biosynthesis |
| ETC | Electron transport chain |
| ID | Inorganic diphosphatase |

**Figure S6:**
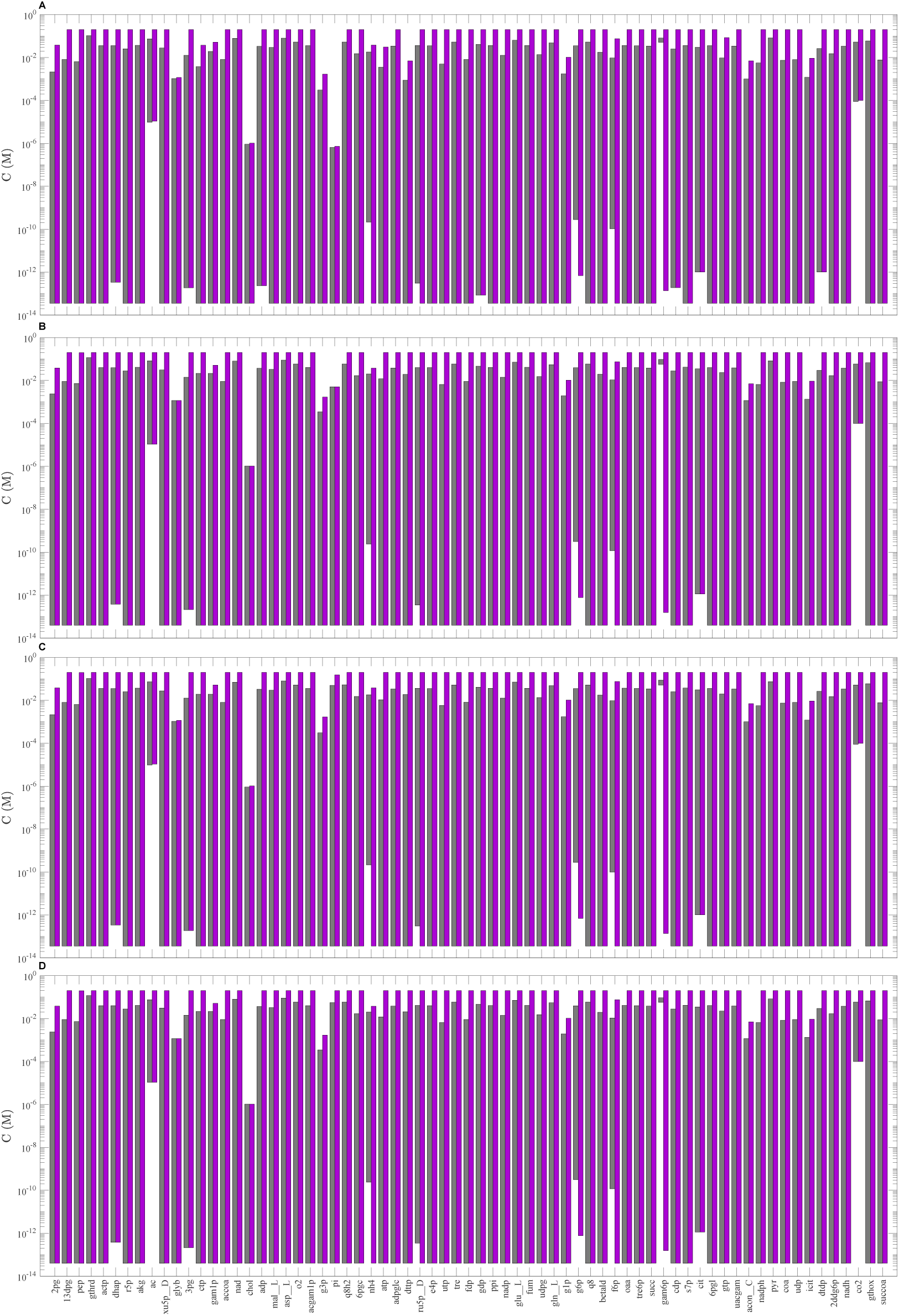
Comparison of feasible concentration ranges in mutants, where either the *Pit* or *Pst* system is active, Related to Fig. 5. (A) *Pit* is the sole phosphate transporter when ⟦pi_p ⟧= 0.01 mM. (B) *Pit* is the sole phosphate transporter when ⟦pi_p⟧ = 69 mM. (C) *Pst* is the sole phosphate transporter when pi p = 0.01 mM. (D) *Pst* is the sole phosphate transporter when ⟦ pi_p⟧ = 69 mM. Color legend is the same as in Fig. 5.

**Figure S7:**
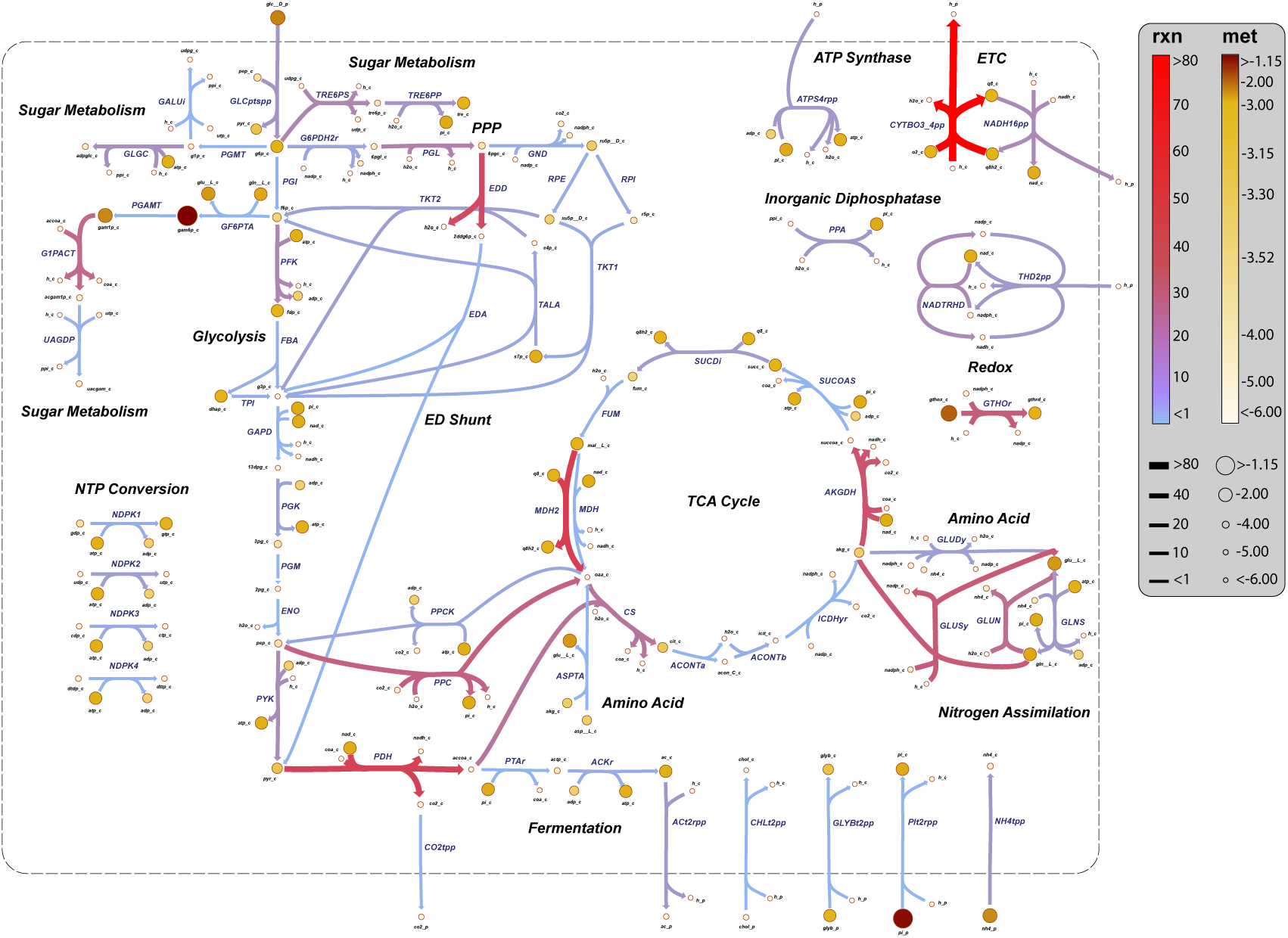
Energy and concentration map, Related to Fig. 3. The description of the legend is identical to Fig. 3A.

**Figure S8:**
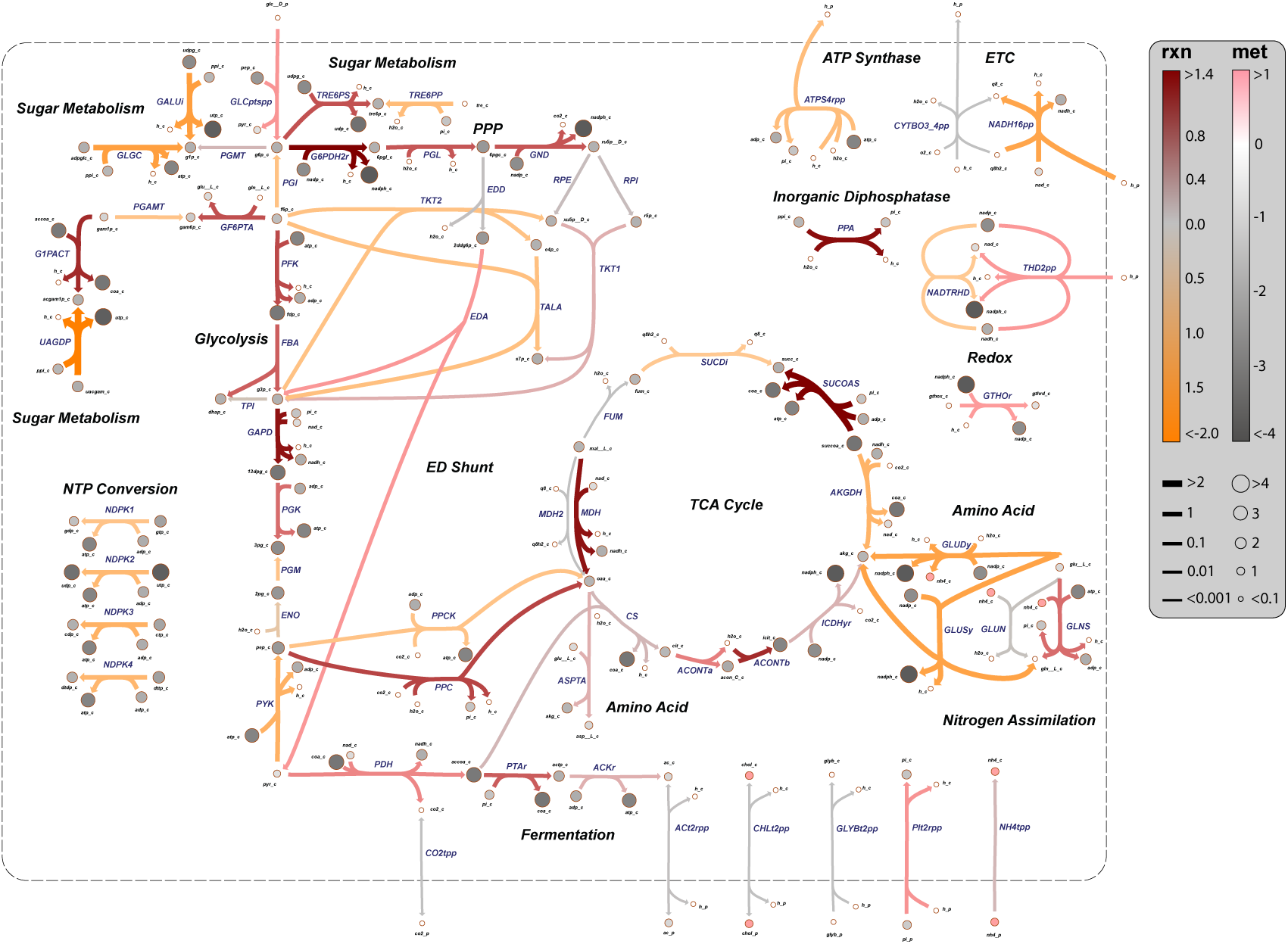
Charge map, Related to Fig. 3. The description of the legend is identical to Fig. 3B.

**Figure S9:**
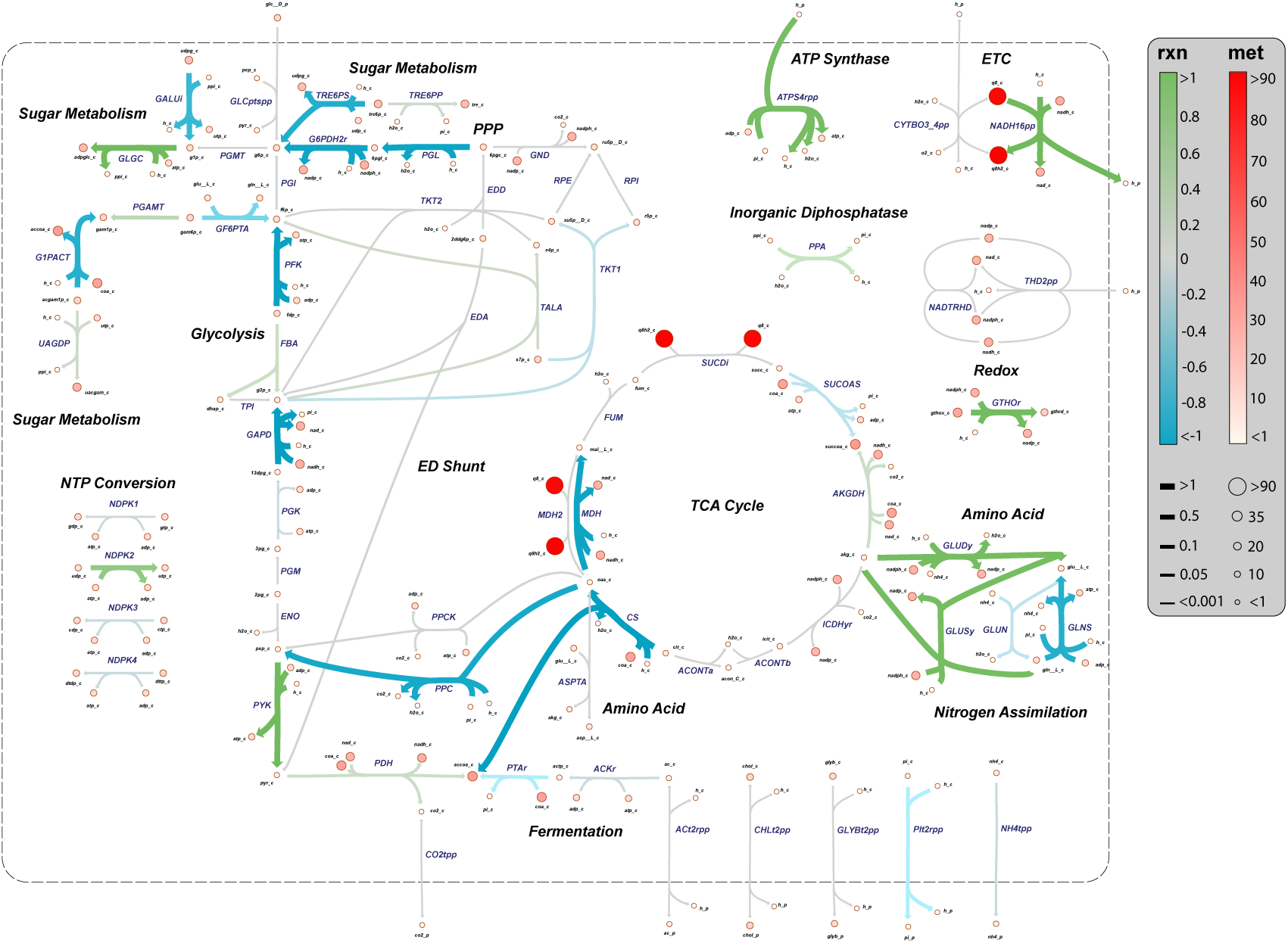
Hydrogen map, Related to Fig. 3. The description of the legend is identical to Fig. 3C.

**Figure S10:**
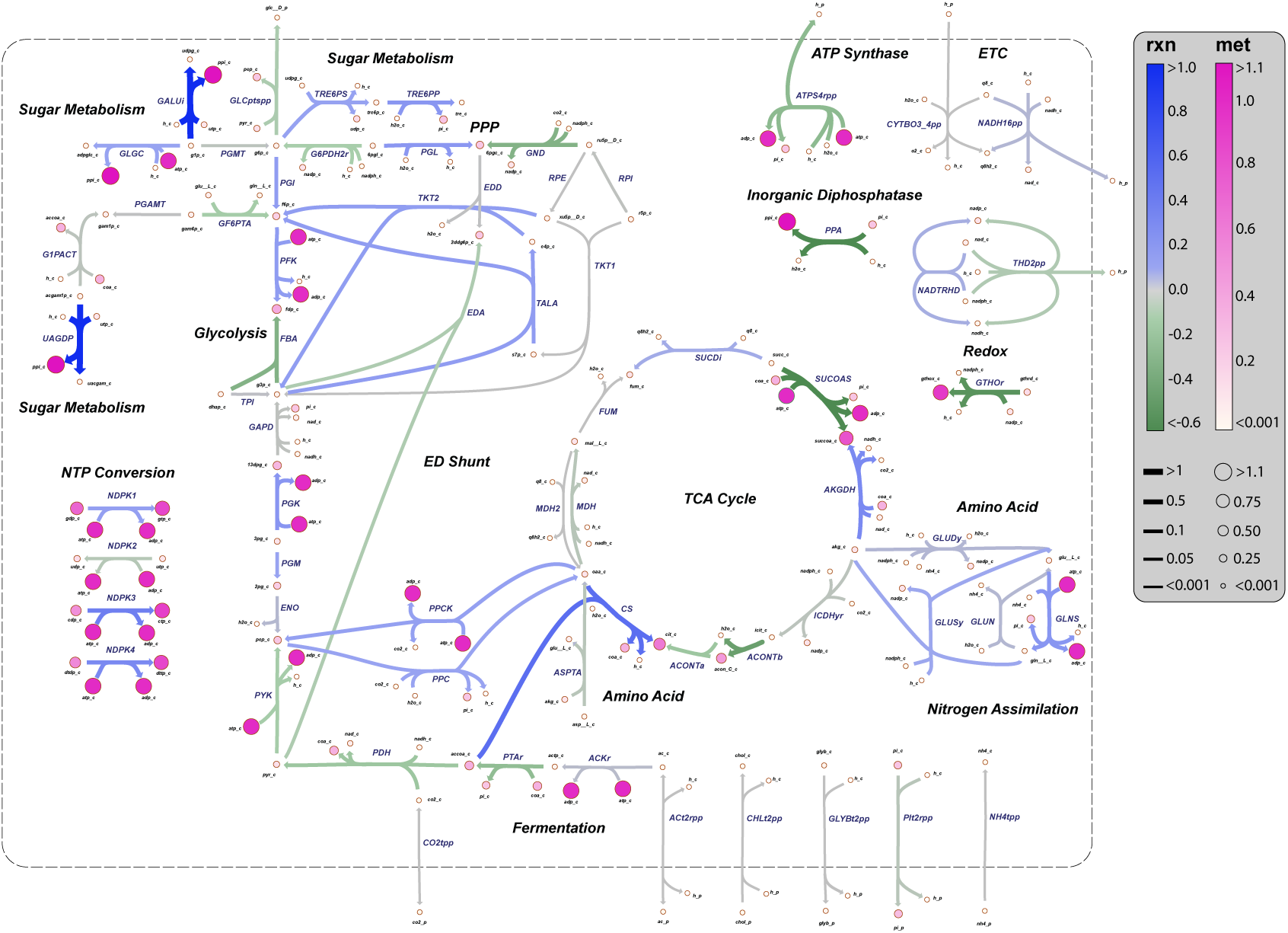
Magnesium map, Related to Fig. 3. The description of the legend is identical to Fig. 3D.

### Global Maps

In this section, we provided enlarged and more detailed representations of the global maps discussed in the main text.

## Notes

### Competing Interest Statement

The authors have declared no competing interest.

## References

1. Alberts, B., Bray, D., Hopkin, K., Johnson, A.D., Lewis, J., Raff, M., Roberts, K., Walter, P.. Essential cell biology. Garland Science; 2013.

2. Kitano, H.. Systems biology: A brief overview. Science 2002;295(5560):1662–1664. doi: 10.1126/science.1069492.

3. Stelling, J.. Mathematical models in microbial systems biology. Curr Opin Microbiol 2004;7(5):513–518. doi: 10.1016/j.mib.2004.08.004.

4. O’Brien, E.J., Monk, J.M., Palsson, B.O.. Using genome-scale models to predict biological capabilities. Cell 2015;161(5):971–987. doi: 10.1016/j.cell.2015.05.019.

5. Palsson, B.O.. Systems biology: Constraint-based reconstruction and analysis. Cambridge University Press; 2015.

6. Orth, J., Thiele, I., Palsson, B.. What is flux balance analysis? Nat Biotechnol 2010;28(3):245–248. doi: 10.1038/nbt.1614.

7. Covert, M.W., Knight, E.M., Reed, J.L., Herrgard, M.J., Palsson, B.O.. Integrating high-throughput and computational data elucidates bacterial networks. Nature 2004;429(6987):92–96. doi: 10.1038/nature02456.

8. Lerman, J.A., Hyduke, D.R., Latif, H., Portnoy, V.A., Lewis, N.E., Orth, J.D., Schrimpe-Rutledge, A.C., Smith, R.D., Adkins, J.N., Zengler, K., Palsson, B.O.. In silico method for modelling metabolism and gene product expression at genome scale. Nat Commun 2012;3(1):1–10. doi: 10.1038/ncomms1928.

9. Maranas, C.D., Zomorrodi, A.R.. Optimization methods in metabolic networks. John Wiley & Sons; 2016.

10. Henry, C.S., Broadbelt, L.J., Hatzimanikatis, V.. Thermodynamics-based metabolic flux analysis. Biophys J 2007;92(5):1792–1805. doi: 10.1529/biophysj.106.093138.

11. Bennett, B.D., Kimball, E.H., Gao, M., Osterhout, R., Van Dien, S.J., Rabinowitz, J.D.. Absolute metabolite concentrations and implied enzyme active site occupancy in Escherichia coli. Nat Chem Biol 2009;5(8):593. doi: 10.1038/nchembio.186.

12. Gerosa, L., van Rijsewijk, B.R.B.H., Christodoulou, D., Kochanowski, K., Schmidt, T.S.B., Noor, E., Sauer, U.. Pseudo-transition analysis identifies the key regulators of dynamic metabolic adaptations from steady-state data. Cell systems 2015;1(4):270–282. doi: 10.1016/j.cels.2015.09.008.

13. Monk, J., Lloyd, C., Brunk, E., Mih, N., Sastry, A., King, Z., Takeuchi, R., Nomura, W., Zhang, Z., Mori, H., Feist, A., Palsson, B.. iml1515, a knowledgebase that computes Escherichia coli traits. Nat Biotechnol 2017;35(10):904–908. doi: 10.1038/nbt.3956.

14. Jeske, L., Placzek, S., Schomburg, I., Chang, A., Schomburg, D.. Brenda in 2019: a european elixir core data resource. Nucleic Acids Res 2019;47(D1):D542–D549. doi: 10.1093/nar/gky1048. URL https://www.brenda-enzymes.org.

15. Volkmer, B., Heinemann, M.. Condition-dependent cell volume and concentration of Escherichia coli to facilitate data conversion for systems biology modeling. PLoS ONE 2011;6(7). doi: 10.1371/journal.pone.0023126.

16. Tepper, N., Noor, E., Amador-Noguez, D., Haraldsdóttir, H., Milo, R., Rabinowitz, J., Liebermeister, W., Shlomi, T.. Steady-state metabolite concentrations reflect a balance between maximizing enzyme efficiency and minimizing total metabolite load. PLoS One 2013;8(9). doi: 10.1371/journal.pone.0075370.

17. Bar-Even, A., Noor, E., Savir, Y., Liebermeister, W., Davidi, D., Tawfik, D.S., Milo, R.. The moderately efficient enzyme: evolutionary and physicochemical trends shaping enzyme parameters. Biochemistry 2011;50(21):4402–4410. doi: 10.1021/bi2002289.

18. Noor, E., Flamholz, A., Bar-Even, A., Davidi, D., Milo, R., Liebermeister, W.. The protein cost of metabolic fluxes: prediction from enzymatic rate laws and cost minimization. PLoS Comput Biol 2016;12(11). doi: 10.1371/journal.pcbi.1005167.

19. Park, J.O., Tanner, L.B., Wei, M.H., Khana, D.B., Jacobson, T.B., Zhang, Z., Rubin, S.A., Hsin-Jung Li, S., Higgins, M.B., Stevenson, D.M., Amador-Noguez, D., Rabinowitz, J.D.. Near-equilibrium glycolysis supports metabolic homeostasis and energy yield. Nat Chem Biol 2019;15(10):1001–1008. doi: 10.1038/s41589-019-0364-9.

20. Chen, X., Alonso, A.P., Allen, D.K., Reed, J.L., Shachar-Hill, Y.. Synergy between ^13^C-metabolic flux analysis and flux balance analysis for understanding metabolic adaption to anaerobiosis in E. coli. Metab Eng 2011;13(1):38–48. doi: 10.1016/j.ymben.2010.11.004.

21. Noor, E., Bar-Even, A., Flamholz, A., Reznik, E., Liebermeister, W., Milo, R.. Pathway thermodynamics highlights kinetic obstacles in central metabolism. PLoS Comput Biol 2014;10(2):e1003483. doi: 10.1371/journal.pcbi.1003483.

22. Alberty, R.. Thermodynamics of biochemical reactions. John Wiley & Sons; 2005.

23. Ly, A., Henderson, J., Lu, A., Culham, D., Wood, J.. Osmoregulatory systems of Escherichia coli: Identification of betainecarnitine-choline transporter family member betu and distributions of betu and trkg among pathogenic and nonpathogenic isolates. J Bacteriol 2004;186(2):296–306. doi: 10.1128/JB.186.2.296-306.2004.

24. Gunasekera, T.S., Csonka, L.N., Paliy, O.. Genome-wide transcriptional responses of Escherichia coli K-12 to continuous osmotic and heat stresses. J Bacteriol 2008;190(10):3712–3720. doi: 10.1128/JB.01990-07.

25. Weber, A., Jung, K.. Profiling early osmostress-dependent gene expression in Escherichia coli using dna macroarrays. J Bacteriol 2002;184(19):5502–5507. doi: 10.1128/JB.184.19.5502-5507.2002.

26. Wood, J.M.. Osmosensing by bacteria: Signals and membrane-based sensors. Microbiol Mol Biol Rev 1999;63(1):230–262. doi: 10.1128/MMBR.63.1.230-262.1999.

27. Seo, S.W., Gao, Y., Kim, D., Szubin, R., Yang, J., Cho, B.K., Palsson, B.O.. Revealing genome-scale transcriptional regulatory landscape of OmpR highlights its expanded regulatory roles under osmotic stress in Escherichia coli K-12 MG1655. Sci Rep 2017;7(1):1–10. doi: 10.1038/s41598-017-02110-7.

28. McLaggan, D., Naprstek, J., Buurman, E., Epstein, W.. Inter-dependence of k+ and glutamate accumulation during osmotic adaptation of Escherichia coli. J Biol Chem 1994;269(3):1911–1917. URL https://www.jbc.org/content/269/3/1911.full.pdf.

29. Schleyer, M., Schmid, R., Bakker, E.P.. Transient, specific and extremely rapid release of osmolytes from growing cells of Escherichia coli K-12 exposed to hypoosmotic shock. Arch Microbiol 1993;160(6):424–431. doi: 10.1007/BF00245302.

30. Sastry, A.V., Gao, Y., Szubin, R., Hefner, Y., Xu, S., Kim, D., Choudhary, K.S., Yang, L., King, Z.A., Palsson, B.O.. The Escherichia coli transcriptome mostly consists of independently regulated modules. Nat Commun 2019;10(1):1–14. doi: 10.1038/s41467-019-13483-w.

31. Chung, H.J., Bang, W., Drake, M.A.. Stress response of Escherichia coli. Compr Rev Food Sci Food Saf 2006;5(3):52–64. doi: 10.1111/j.1541-4337.2006.00002.x.

32. Seo, S.W., Kim, D., O’Brien, E.J., Szubin, R., Palsson, B.O.. Decoding genome-wide GadEWX-transcriptional regulatory networks reveals multifaceted cellular responses to acid stress in Escherichia coli. Nat Commun 2015;6(1):1–8. doi: 10.1038/ncomms8970.

33. Kannan, G., Wilks, J.C., Fitzgerald, D.M., Jones, B.D., BonDurant, S.S., Slonczewski, J.L.. Rapid acid treatment of Escherichia coli: transcriptomic response and recovery. BMC Microbiol 2008;8(1):37. doi: 10.1186/1471-2180-8-37.

34. Lane, N.. The vital question: energy, evolution, and the origins of complex life. WW Norton & Company; 2015.

35. Mulkidjanian, A., Galperin, M., Koonin, E.. Co-evolution of primordial membranes and membrane proteins. Trends Biochem Sci 2009;34(4):206–215. doi: 10.1016/j.tibs.2009.01.005.

36. Willsky, G.R., Malamy, M.H.. Characterization of two genetically separable inorganic phosphate transport systems in Escherichia coli. J Bacteriol 1980;144(1):356–365. URL https://www.ncbi.nlm.nih.gov/pmc/articles/PMC294655/pdf/jbacter00571-0370.pdf.

37. Nesmeyanova, M.. Polyphosphates and enzymes of polyphosphate metabolism in Escherichia coli. Biochemistry 2000;65(3):309–314. URL http://protein.bio.msu.ru/biokhimiya/contents/v65/pdf/bcm_0309.pdf.

38. McCleary, W.. Molecular mechanisms of phosphate homeostasis in Escherichia coli; chap. 17. InTech; 2017:333–357.

39. Jacob, F.. Evolution and tinkering. Science 1977;196(4295):1161–1166. doi: 10.1126/science.860134.

40. Schuetz, R., Zamboni, N., Zampieri, M., Heinemann, M., Sauer, U.. Multidimensional optimality of microbial metabolism. Science 2012;336(6081):601–604. doi: 10.1126/science.1216882.

41. Dyson, F.. Origins of life. Cambridge University Press; 1999.

42. Brack, A.. The molecular origins of life: Assembling pieces of the puzzle. Cambridge University Press; 1998.

43. Berg, J.M., Tymoczko, J.L., Stryer, L.. Biochemistry. Fifth ed.; Freeman: New York; 2002.

44. Urbansky, E.T., Schock, M.R.. Understanding, deriving, and computing buffer capacity. J Chem Educ 2000;77(12):1640. doi: 10.1021/ed077p1640.

45. Prausnitz, J., Lichtenthaler, R., De Azevedo, E.. Molecular thermodynamics of fluid-phase equilibria. Pearson Education; 1998.

46. Pitzer, K., Kim, J.. Thermodynamics of electrolytes. IV. Activity and osmotic coefficients for mixed electrolytes. J Am Chem Soc 1974;96(18). doi: 10.1021/ja00825a004.

47. Goldberg, R.N., Tewari, Y.B.. Thermodynamics of the disproportionation of adenosine 5-diphosphate to adenosine 5-triphosphate and adenosine 5-monophosphate: I. equilibrium model. Biophys Chem 1991;40(3):241–261. doi: 10.1016/0301-4622(91)80025-m.

48. Flamholz, A., Noor, E., Bar-Even, A., Milo, R.. equilibrator—the biochemical thermodynamics calculator. Nucleic Acids Res 2012;40(D1):D770–D775. doi: 10.1093/nar/gkr874.

49. Steuer, R.. Computational approaches to the topology, stability and dynamics of metabolic networks. Phytochemistry 2007;68(16-18):2139–2151. doi: 10.1016/j.phytochem.2007.04.041.

50. Palsson, B.Ø.. Systems biology: Simulation of dynamic network states. Cambridge University Press; 2011.

51. Lengyel, S.. Deduction of the Guldberg–Waage mass action law from Gyarmati’s governing principle of dissipative processes. J Chem Phys 1988;88(3):1617–1621. doi: 10.1063/1.454140.

52. Goldbeter, A.. Biochemical oscillations and cellular rhythms: the molecular bases of periodic and chaotic behaviour. Cambridge University Press; 1997.

53. Kay, A.R., Blaustein, M.P.. Evolution of our understanding of cell volume regulation by the pump-leak mechanism. J Gen Physiol 2019;151(4):407–416. doi: 10.1085/jgp.201812274.

54. Liu, B., Poolman, B., Boersma, A.. Ionic strength sensing in living cells. ACS Chem Biol 2017;12(10):2510–2514. doi: 10.1021/acschembio.7b00348.

55. Slonczewski, J.L., Fujisawa, M., Dopson, M., Krulwich, T.A.. Cytoplasmic pH measurement and homeostasis in bacteria and archaea. Adv Microb Physiol 2009;55:1–317. doi: 10.1016/s0065-2911(09)05501-5.

56. Stein, W.. Transport and diffusion across cell membranes. Elsevier; 2012.

57. Sambrook, J.. Molecular cloning: A laboratory manual; vol. 999. Cold Spring Harb Lab Press Cold Spring Harb NY; 2001.

58. Noor, E., Flamholz, A., Liebermeister, W., Bar-Even, A., Milo, R.. A note on the kinetics of enzyme action: a decomposition that highlights thermodynamic effects. FEBS Lett 2013;587(17):2772–2777. doi: 10.1016/j.febslet.2013.07.028.

59. Schultz, S.G., Solomon, A.K.. Cation transport in Escherichia coli: I. intracellular Na and K concentrations and net cation movement. J Gen Physiol 1961;45(2):355–369. doi: 10.1085/jgp.45.2.355.

60. Schultz, S.G., Wilson, N.L., Epstein, W.. Cation transport in Escherichia coli: II. intracellular chloride concentration. J Gen Physiol 1962;46(1):159–166. doi: 10.1085/jgp.46.1.159.

61. Ivancevic, V.G., Ivancevic, T.T.. Applied differential geometry: A modern introduction. World Scientific; 2007.

62. Lee, J.M.. Riemannian manifolds: An introduction to curvature; vol. 176. Springer Science & Business Media; 2006.

63. Boyd, S.P., Vandenberghe, L.. Convex optimization. Cambridge university press; 2004.

64. Bussieck, M.R., Meeraus, A.. General algebraic modeling system (GAMS). In: Modeling languages in mathematical optimization. Springer; 2004:137–157. doi: 10.1007/978-1-4613-0215-5_8.

65. Tawarmalani, M., Sahinidis, N.V.. A polyhedral branch-and-cut approach to global optimization. Math Program 2005;103(2):225–249. doi: 10.1007/s10107-005-0581-8.

